# SIRT2 protects against Japanese encephalitis virus infection in mice

**DOI:** 10.1101/2025.03.24.645101

**Authors:** Perumal Arumugam Desingu, Lavanya Dindi, Krishnega Murugasamy, Ankit Kumar Tamta, Venketsubbu Ramasubbu, Sukanya Raghu, Amarjeet Shrama, Raju S. Rajmani, Nagalingam R. Sundaresan

**Affiliations:** Department of Microbiology and Cell Biology, Indian Institute of Science, Bengaluru 560012, India; Department of Virus Epidemiology, Vector Dynamics & Public Health, Institute of Advanced Virology, Bio 360 Life Sciences Park, Thonnakkal, Trivandrum, Kerala-695 317, India; Centre for Infectious Disease Research, Indian Institute of Science, Bengaluru 560012, India

**Keywords:** SIRT2, JEV, Flavivirus, NF-κB, deacetylation, Beclin-1, autophagy

## Abstract

Japanese encephalitis virus (JEV) is a mosquito-borne zoonotic RNA virus that causes Japanese encephalitis (JE) and poses a major threat to public health in Southeast Asia and the Western Pacific. Current strategies rely on prophylactic methods to prevent disease, as no effective antiviral therapy exists. Here, we report that SIRT2, an NAD+-dependent deacetylase enzyme, mediates antiviral activity against JEV infection in mice. Interestingly, our study reveals that SIRT2 is downregulated in JEV infection, SIRT2 genetic deficiency/small molecule inhibition increases viral yield in neuronal cells and mice brains thereby reducing the survival rate in the infected mice, whereas SIRT2 gene therapy to the JEV-infected mice by Adeno-associated virus vector reduced the JEV load in mice brains and improved the survival rate. SIRT2 deficiency activates inflammatory cytokines and chemokines response in the JEV-infected mice brains through activating NF-κB transcription factor. Mechanistically, SIRT2 deacetylates NF-κB to reduce the transcriptional factor activity of NF-κB that down-regulates the Beclin-1-mediated autophagy, which is needed for the JEV replication. Overall, the present findings establish SIRT2 as a potential regulator of JEV infection.

## Introduction

JE is a vector-borne zoonotic viral disease caused by the Japanese encephalitis virus (JEV)^1–3^. JEV exhibits close resemblance to other members of the Flaviviridae family, including dengue virus, Zika virus, yellow fever virus, and West Nile virus^1–4^. Estimates suggest that JE is endemic in about twenty-four countries in the Southeast Asia, South Asia, and Western Pacific geographic regions. Almost 68 thousand clinical cases occur every year, posing a risk to three billion people^1–3,5^. In the endemic areas, children aged 2 to 15 years are more susceptible to JEV, and exhibit adverse outcomes, with a 30% fatality rate and permanent neurologic or psychiatric sequelae of 30-50% in encephalitis patients^1–3,5^. Currently, vaccine administration in the endemic area is the only intervention against JEV^1^. The absence of clinically available therapeutics raises serious health threats and warrants the identification of master regulators of host proteins in the virus pathogenesis to determine an antiviral against JEV. Because of the high rate of mutations in RNA viruses^6–9^, it would be appropriate to target host proteins and the evolutionarily conserved host mechanisms for therapeutic intervention.

In this line, we recently identified the inhibition of NAD+-dependent host enzyme, poly (ADP-ribose) polymerases (PARP1) inhibition, as a potential antiviral strategy^1^. Possibly other NAD+-dependent host enzymes may also play a role in the JEV infection and the identification of such enzymes may be useful in designing a host protein-directed antiviral against JEV infection. In this context, sirtuins (SIRTs) are well-studied NAD+-dependent enzymes that play roles in ageing, metabolism, and cell survival, and they have been examined in relation to viral infections recently. Sirtuins are class III histone deacetylases that depend on NAD^+^ as a cofactor for their enzymatic activity. Mammals have seven sirtuins (SIRT1-SIRT7), ubiquitously expressed with distinct sub-cellular localisation^10,11^. SIRTs govern several cellular processes through multiple targets and have critical implications in health and diseases^12–14^. Of them, the major cytosolic protein SIRT2 is known to regulate cell division, inflammation, autophagy, mitophagy, oxidative stress, and senescence^11,15,16^. It has been shown that SIRT1-SIRT7 knockdown enhances RNA and DNA virus replication in cell culture models^14,17,18^. In contrast, treatment with the SIRT2 inhibitor AGK2 was shown to inhibit the replication of DNA viruses in mouse models^19,20^. Recent reports indicate that SIRT2 protects aged mice from severe COVID-19 by suppressing age-related cGAS activation and inflammation^21^. In this context, understanding the role of SIRT2 in distinct viral pathology, especially encephalitis-causing RNA viruses in mice models, and exploring molecular mechanisms in the evolutionarily conserved signalling axis, holds direct therapeutic relevance for preclinical studies. Further, this knowledge can be extended to develop therapeutics against several other viruses that utilise the same host proteins and similar mechanisms for their pathogenesis.

Several RNA viruses including the hepatitis C virus (HCV)^22,23^, porcine reproductive and respiratory syndrome virus (PRRSV)^24,25^, Severe acute respiratory syndrome coronavirus (SARS-CoV)^26^, influenza A virus (IAV)^27,28^, JEV^1^, Dengue virus (DENV)^29,30^, Zika virus (ZIKV)^31,32^, Chikungunya virus (CHIKV)^33^, Human rhinovirus (HRV)^34^, Enterovirus 71 (EV-A71)^35^, poliovirus^36,37^, Coxsackievirus ^38,39^, encephalomyocarditis virus (EMCV)^40^, Foot-and-mouth disease virus (FMDV)^41^, Newcastle disease virus (NDV)^42^ etc., exploit evolutionarily conserved cellular autophagy for viral replication by different molecular mechanisms. In this context, the functional role of SIRT2 in RNA viruses-induced autophagy has not yet been explored. Recent studies highlight the role of SIRT2 in inflammatory signalling mediated by the NLRP3 inflammasome^43,44^, as well as its involvement in age-associated inflammation^45^ and COVID-19-induced inflammation in older mice^21^. Of note, different RNA viruses target the host protein nuclear factor-κB (NF-κB), to modulate the inflammatory cytokine and chemokine transcription (reviewed in^46–49^). The role of phosphorylation of IkB in the NF-κB-IkB axis^50,51^ has been well-studied in viral infections. In contrast, the role of post-translational modifications, particularly acetylation of NF-κB, on its transcriptional factor activity-mediated regulation of RNA virus-induced autophagy remains unexplored.

In our study, we uncover a key role for SIRT2 in regulating JEV replication and improving the survival in JEV-infected mice. Our *in vitro* and *in vivo* model systems demonstrate that this effect is SIRT2-mediated deacetylation of NF-κB thereby regulating virus-induced autophagy and inflammatory cytokine. Further, we report that SIRT2 mediated NF-κB regulation negatively affects the virus-induced autophagy, which is needed for virus replication. Thereby SIRT2 exhibits an antiviral role in neuronal cells and mice infected with JEV.

## Results

### The JEV infection down-regulates SIRT2 in the *in vitro* and *in vivo* models

Our recent study determined the role of NAD^+^-dependent enzyme PARP1 in JEV replication^1^. However, the role of other NAD+-dependent enzymes remains unexplored. We were interested in exploring the role of NAD^+^-dependent deacetylase enzymes, Sirtuins ^52^, in JEV replication. Because JEV replicates in the cytoplasm^1–3,5^, we wanted to understand the importance of SIRT2, a cytoplasmic sirtuin^10,11^, in the regulation of JEV replication. To begin with, we aimed to identify the molecular mechanisms occurring at the early viral replication time point that can be targeted as an antiviral against JEV infection. To determine the same, we infected the neuro2a cells with one multiplicity of infection (MOI) of JEV and observed the viral protein levels and infective virus yield over time. We found that the maximum rate of JEV multiplication occurs between 24-36 hours post-infection (hpi) (**Figure 1A-1B**) and therefore considered 30 hpi as an early accelerating phase of viral replication (**Figure 1B**). At this time point, SIRT2 was significantly down-regulated (**Figure 1C**).

**Figure 1.**
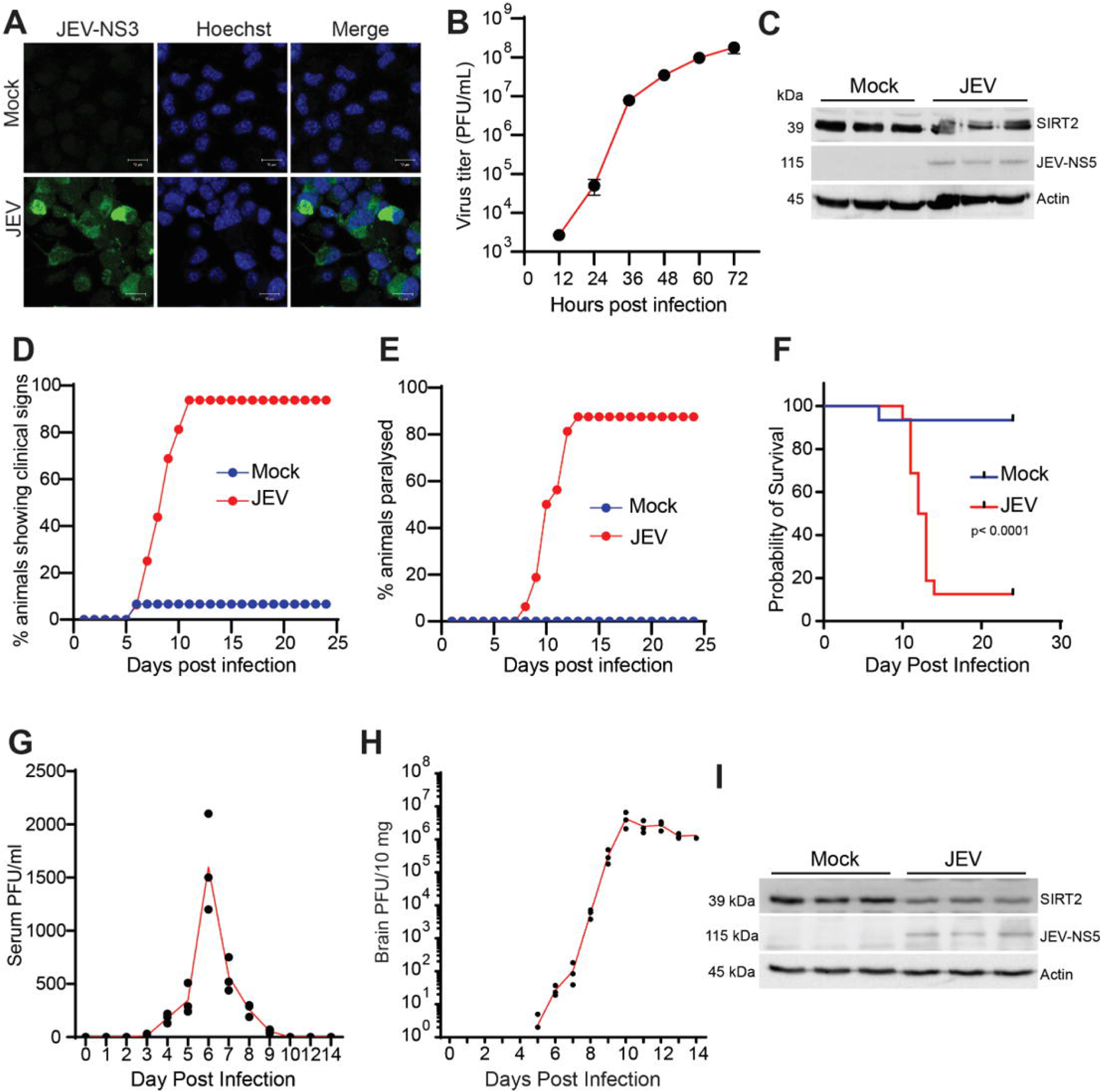
SIRT2 downregulated in neuro2a cells and mice brains upon JEV infection. (**A**) The representative confocal images of JE virus (one MOI) or mock-infected neuro2a cells at 30 hpi; JEV-NS3 protein stained in green, and the nuclei stained by Hoechst 33342 in blue. The scale bar =10 mM. (**B**) The virus load upon 1 MOI of JE virus infection in neuro2a cells over time expressed in plaque-forming units (PFUs). n=3 independent experiments; each experiment performed by duplicates. (**C**) Representative immunoblot image displaying a reduction in SIRT2 levels in neuro2a cells upon 1 MOI of JE virus infection at 30 hpi. The JEV protein NS5 confirms JEV infection in the cells. n=3 independent experiments; each experiment performed by duplicates. (**D-I**) 10-day-old mice were infected through the JE virus of 10^5^ PFU by the foot-pad route of inoculation; n=15-16 mice per group. Graphical representation depicting the percentage of mock or JE virus-infected mice expressing clinical signs (**D**) and paralysis (**E**) at various dpi. (**F**) The percentage of mice surviving upon JEV mice at various dpi up to 24^th^ dpi, in Kaplan-Meier survival percentage curve (n=15-16 mice each group). The p-value was calculated using the Log-rank (Mantel-Cox) test. Graphically representing the JEV load (PFUs) in the serum (**G**) and brain (**H**) of wild-type mice at various dpi; n=3 mice per dpi. (**I**) Representative immunoblot indicating the reduction in SIRT2 levels in the mice brain upon JEV-infection on 8^th^ dpi. The detection of NS5 confirms the virus infection. n=6 mice in each group.

Further, to investigate the role of SIRT2 in an *in vivo* system, we studied the brains of JEV-infected mice. Since JEV infection is spread through mosquito bites and causes encephalitis in young children^53^, we infected 10 days old mice pups with 10^5^ plaque-forming units (PFU) of JEV by foot-pad route to mimic natural infection. The infected mice were monitored for the display of clinical signs, paralysis, and survival every 24 hours post-infection. The standard clinical signs observed include fluffy coat, ruffled fur, reluctance to move, hunchback posture, circling and curling behaviour, ataxia, and tremors on the 6^th^ day post-infection (dpi). By the 11th dpi, we observed that about 90 per cent of animals displayed clinical signs (**Figure 1D**). The mice infected with JEV started showing paralysis at the 8^th^ dpi, which peaked by the 13^th^ day (**Figure 1E**). We observed mortality in JEV-infected mice 10^th^ dpi onwards (**Figure 1F**). The serum viral load was detectable 3^rd^ dpi onwards and peaked on the 6^th^ dpi, coinciding with the onset of clinical signs (**Figure 1D, 1G**). Interestingly, serum viral load started declining when the virus started multiplying in the mice’s brain (**Figure 1G, 1H**). The maximum rate of viral replication in the brain, primary target site of JEV, occurred between 7 and 9^th^ dpi (**Figure 1H**). Thus, 8^th^ dpi was noted to be an early accelerating phase of viral replication in the brain of mice (**Figure 1H**). Next, we measured SIRT2 levels in mock- or JEV-infected mouse brains on 8^th^ dpi. In line with our *in vitro* findings, SIRT2 levels were significantly downregulated at 8^th^ dpi in JEV-infected mouse brains (**Figure 1I**). Thus, significant SIRT2 downregulation coincides with the early accelerating phase of viral replication in both *in vitro* and *in vivo* study systems.

### SIRT2 deficiency enhances JEV replication in both *in vitro* and *in vivo* models

Since SIRT2 is downregulated in JEV infection, we infected SIRT2-depleted neuro2a cells with 1 MOI JEV to study the role of SIRT2 in viral multiplication. We found that viral titre was significantly high in SIRT2 knockdown conditions (**Figure 2A, 2B**). In contrast, SIRT2 overexpression remarkably reduced viral titre in neuro2a cells (**Figure S1A, S1B**), thus suggesting a role for SIRT2 in JEV multiplication. To validate our findings in the *in vivo* model system, we infected SIRT2 knock-out (KO) mice with JEV through the footpad route (**Figure 2C**). We observed that the viral PFU levels in serum reached a maximum in SIRT2-KO mice earlier than their WT counterparts (**Figure 2D-2E**). Similarly, the viral load in the JEV-infected SIRT2-KO mice brain was much higher than in WT (**Figure 2F**). Likewise, clinical signs and paralysis appeared earlier in SIRT2-KO mice than WT mice. The SIRT2-KO group also showed an increased percentage of animals exhibiting clinical signs and paralysis (**Figure 2G-2H**). Further, all the SIRT2-KO mice infected with JEV died by the 14^th^ dpi, with significantly reduced survival of JEV-infected SIRT2-KO mice (**Figure 2I**). To ensure that the observed pathological signs are due to JEV infection in the brain rather than serum viral load, we employed the intracerebral route of JEV infection **(Figure 2J)**. In corroboration with our findings from the foot-pad route of viral inoculation, the intracerebral route of viral infection also rendered SIRT2-KO mice more susceptible to JEV pathogenesis as indicated by increased viral titre, earlier onset of clinical sign, paralysis, and reduced survival rate (**Figure 2K-2N**). Since even WT mice exhibited very severe disease at this viral dose and clinical outcome differences between WT and SIRT2-KO mice were modest, we conducted an experiment using a sub-lethal viral dose of 5 × 10^2 PFU in 14-day-old WT and SIRT2-KO mice to evaluate clinical outcomes. We observed a significant reduction in survival rates among the SIRT2-KO mice (**Figure 2O**). Thus, SIRT2 deficiency in the *in vitro* and *in vivo* models enhances JE virus replication thereby reducing the survival of JEV-infected mice, and SIRT2 overexpression helps curtail JEV titre *in vitro*.

**Figure 2.**
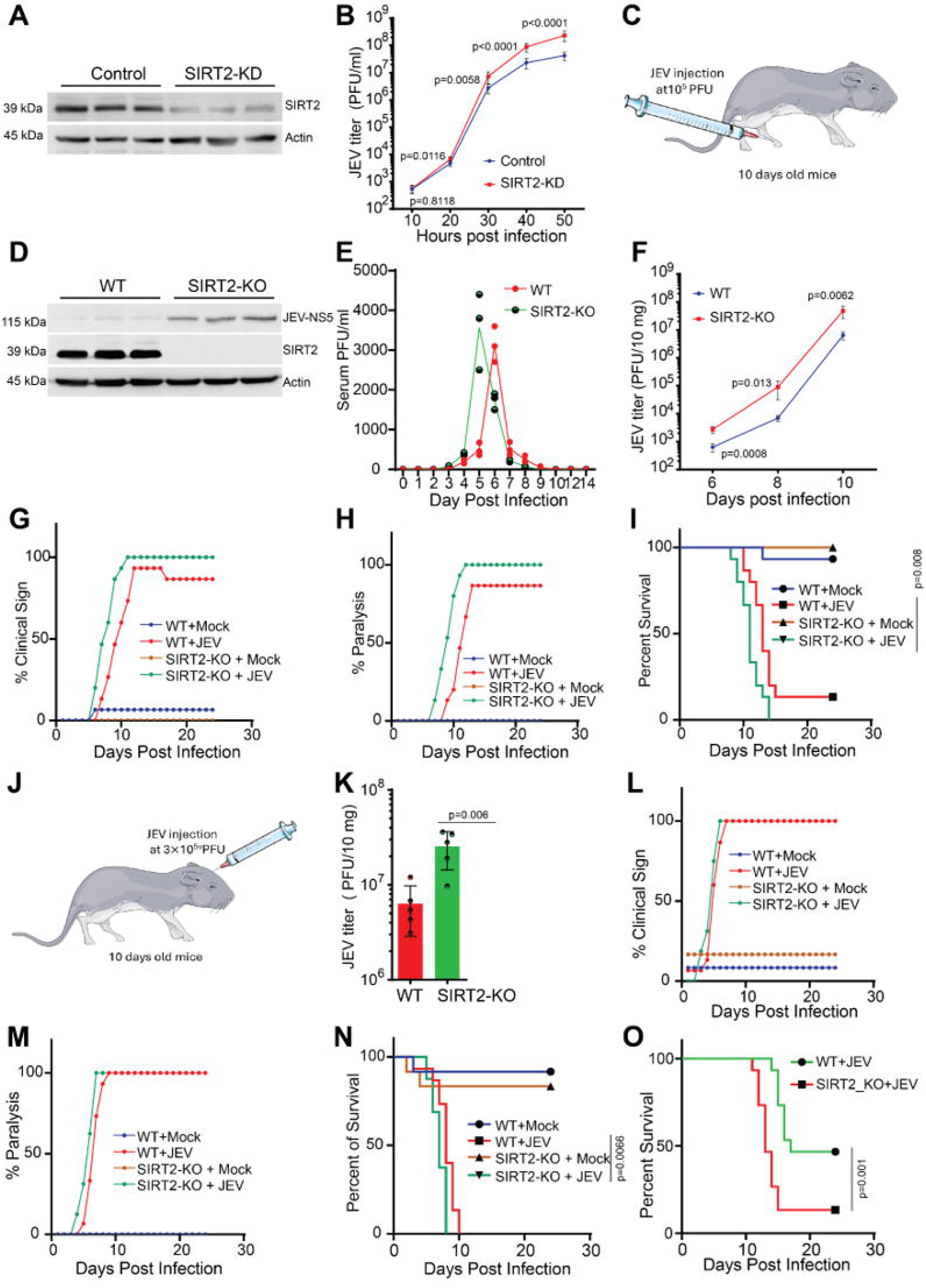
SIRT2 genetic deficiency increases JEV multiplication in neuro2a cells and mice brains. (**A**) Representative western blotting confirms SIRT2 knock-down in neuro2a cells by siRNA. (**B**) Graph displaying the viral PFU levels at the various hpi of control or SIRT2 knock-down neuro2a cells infected with JE-Virus. n=3 independent experiments; triplicates were used in each experiment. P values are calculated using Student’s t-test. (**C-I**) 10-day-old mice were infected through the JE virus of 10^5^ PFU by the foot-pad route of inoculation. (**C**) Schematic representation of JEV administration in mice by foot-pad route. (**D**) Representative immuno-blotting validating the absence of SIRT2 in the SIRT2^-/-^ (SIRT2- KO) mice brain; JE virus infection in the mice brain was confirmed by detecting JEV-NS5 protein. (**E**) The graph represents JE virus PFUs at various dpi in the serum of WT or SIRT2-KO mice infected with JE virus. n=3 mice per dpi. (**F**) The graph represents JE virus PFUs at various dpi in the brains of JE virus-infected WT or SIRT2-/- mice. n=5 mice per dpi. P values are calculated using Student’s t-test. (**G**) The graph represents the percentage of animals expressing clinical signs at various dpi in JE virus-infected WT or SIRT2^-/-^ mice (n=15 mice per group). (**H**) The graph depicts the percentage of animals displaying paralysis in JE virus-infected WT or SIRT2^-/-^ mice at various dpi (n=15 mice in each group). (**I**) The percentage of animals surviving in JE virus-infected WT or SIRT2^-/-^ mice at various dpi up to 24th dpi, in the Kaplan-Meier survival percentage curve. n=15 mice in each group; the p-value was calculated using the Log-rank (Mantel-Cox) test. (**J-N**) 10-day-old mice were inoculated with JE virus (3x10^3^ PFUs) by an intra-cerebral route of inoculations. (**J**) Schematic representation of JEV administration in mice by an intra-cerebral route. **(K)** The graph represents JEV PFUs at 6^th^ dpi in WT or SIRT2^-/-^ mice brains. n=5 mice per group; the p-value was calculated using Student’s t-test. (**L**) The graph represents the percentage of animals expressing clinical signs at various dpi in JE virus-infected WT or SIRT2^-/-^ mice. n=12-16 mice in each group. (**M**) The graph depicts the percentage of animals displaying clinical paralysis at various dpi in JE virus-infected WT or SIRT2^-/-^ mice (n=12-16 mice in each group). (**N**) The percentage of animals surviving at various dpi up to 24th dpi in JE virus-infected WT or SIRT2^-/-^ mice in the Kaplan-Meier survival percentage curve. n=12-16 mice in each group; the p-value was calculated using the Log-rank (Mantel-Cox) test. (**O**) 14-day-old WT and SIRT2-KO mice were infected with the JE virus at 5 × 10^2 PFU by the foot-pad route of inoculation. The percentage of animals surviving at various dpi up to 24th dpi in JE virus-infected WT or SIRT2^-/-^ mice in the Kaplan-Meier survival percentage curve. n=15 mice in each group; the p-value was calculated using the Log-rank (Mantel-Cox) test.

### Pharmacological inhibition of SIRT2 by AGK2 enhances JEV replication

Previous studies have shown that SIRT1-SIRT7 knockdown enhances a few RNA and DNA virus replication in cell culture models^14,17,18^. In contrast, treatment with the SIRT2 inhibitor, AGK2 was shown to inhibit the replication of DNA viruses in mice models^19,20^. Conversely, our study suggests that SIRT2 depletion enhances viral replication in mouse models. We next asked if SIRT2 inhibition, using the small molecule inhibitor AGK2, can enhance JEV replication. We treated neuro2a cells with AGK2 and observed significantly increased viral titers (**Figure 3A**). Further, we tested the effect of AGK2 in JEV infected mice. We found that JEV-infected mice, when treated with AGK2, showed early incidence and an increase in the percentage of clinical signs and paralysis (**Figure 3B, 3C**). AGK2 treatment also reduced the survival rate in JEV-infected mice (**Figure 3D**). We also observed an early appearance and increased serum viral load upon AGK2 treatment (**Figure 3E**). Similarly, the viral load in mice brain was elevated upon AGK 2 treatment (**Figure 3F, 3G**). Consistently, in mice infected via the intracerebral route, AGK2 treatment resulted in an early onset of clinical signs and paralysis (**Figure 3H, 3I**) and a reduction in survival rate compared to their mock infected counterparts (**Figure 3J**). Next, to evaluate the effect of AGK2 on other SIRTs, we administered AGK2 to JEV-infected WT and SIRT2-KO mice and observed that the differences in survival rates between JEV-infected wild-type mice treated with AGK2 and JEV-infected SIRT2-KO mice treated with AGK2 were statistically insignificant (**Figure 3K**). Thus, SIRT2 inhibition using AGK2 exacerbated the clinical outcomes of JE in mice.

**Figure 3.**
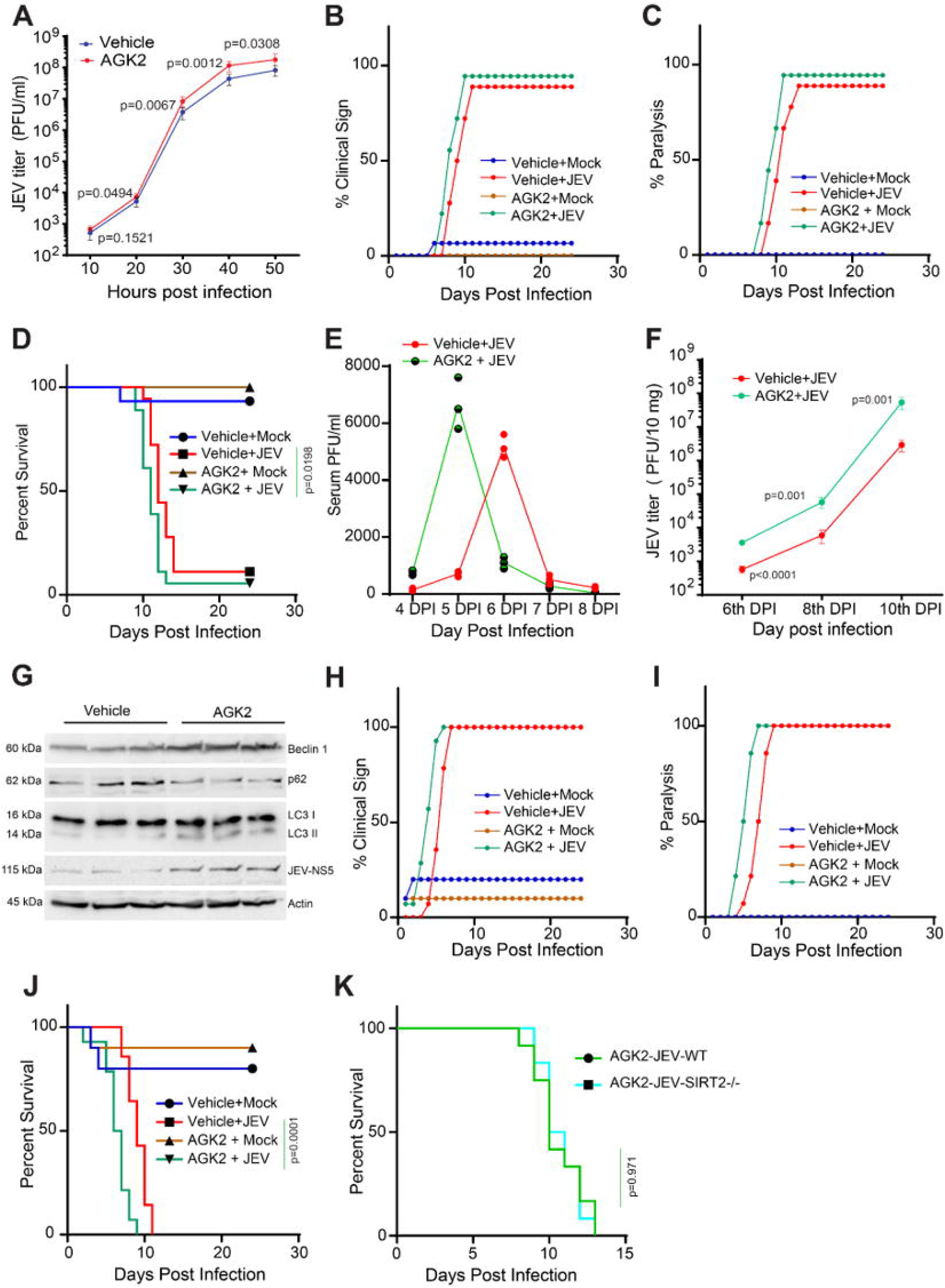
AGK2 treatment exacerbated clinical manifestation in JEV-infected mice through induction of autophagy. **(A)** Graph displaying the viral PFU levels at the various hpi of Vehicle or AGK2 treated neuro2a cells infected with JE-Virus. n=3 independent experiments; triplicates were used in each experiment. P values are calculated using Student’s t-test. (**B-F**) 10-day-old WT mice were inoculated with JE virus of 10^5^ PFUs by the foot-pad route; AGK2 treatment started from 1^st^ dpi onwards for every day up to 24^th^ dpi through the intraperitoneal (IP) route. (**B**) Graph representing the percent animals expressing clinical signs at various dpi in vehicle or AGK2 treated JE virus infected mice; n=15-18 mice per group. (**C**) Graph depicting the percent of animals displaying paralysis at various dpi in JE virus-infected mice upon vehicle or AGK2 treatment. n=15-18 mice per group. (**D**) The percentage of mice surviving after JE virus infection upon vehicle or AGK2 treatment at various dpi is depicted in the Kaplan-Meier survival curve. Statistical analysis was done using Log rank (Mantel-Cox) test; n=15-18 mice per group. Graph representing JE virus load in **(E)** serum (PFU/ml) (n=3 mice per dpi) and **(F)** brain (PFU/10mg) (n=5 mice per dpi), upon vehicle or AGK2 treatment in JE virus-infected mice at various dpi. The p-value was calculated using Student’s t-test. (**G**) Representative immunoblot displaying differences in autophagy markers in the brains of the vehicle or AGK2 treated mice at 8^th^ dpi of JE virus infection (n=3 mice each group). (**H-J**) 10-day-old Wild-Type mice were inoculated with 3x10^3^ PFUs of JEV virus by an intra-cerebral route of inoculation, and AGK2 treatment started from first dpi for every day up to 24^th^ dpi by the IP route of inoculation; n=10-14 mice each group. Graph displaying the percent of animals showing **(H)** clinical signs and **(I)** paralysis, at various dpi in JE virus-infected mice upon vehicle or AGK2 treatment. (n=10-14 mice each group). (**J**) The survival of JE virus-infected mice treated with vehicle or AGK2 up to 24th dpi in the Kaplan-Meier survival percentage curve. The p-value was calculated using the Log-rank (Mantel-Cox) test. (n=10-14 mice per group). (**K**) 10-day-old WT or SIRT2-KO mice were inoculated with JE virus of 10^5^ PFUs by the foot-pad route; AGK2 treatment started from 1^st^ dpi onwards for every day up to 24^th^ dpi through the intraperitoneal (IP) route. The percentage of mice surviving in the JEV-infected WT or JEV-infected SIRT2-KO mice upon AGK2 treatment at various dpi is depicted in the Kaplan-Meier survival curve. Statistical analysis was done using Log rank (Mantel-Cox) test; n=12 mice per group.

### SIRT2 gene therapy protects against JEV infection in mice

Next, we were interested in exploring the therapeutic efficacy of SIRT2 gene therapy against JEV infection in mice. For this, we overexpressed SIRT2 in mock or JEV-infected mice brains using the Adeno-associated virus (AAV) system (**Figure 4A**). Interestingly, AAV-rSIRT2 treatment after JEV infection had significantly less viral load as compared to AAV-Null treated, JEV infected mice brains (**Figure 4A, 4B**). In corroboration with reduced viral load in the JEV-infected WT mice brains, we also observed that SIRT2 gene therapy significantly reduced clinical signs and paralysis and enhanced survival rates in the JEV-infected wild-type mice compared to control AAV-vector treated JEV-infected mice (**Figure 4C-4E**). Overall, our results suggest that SIRT2 is a potential regulator of JEV infection in mice, and it could be used as a host protein-directed antiviral against JEV infection.

**Figure 4.**
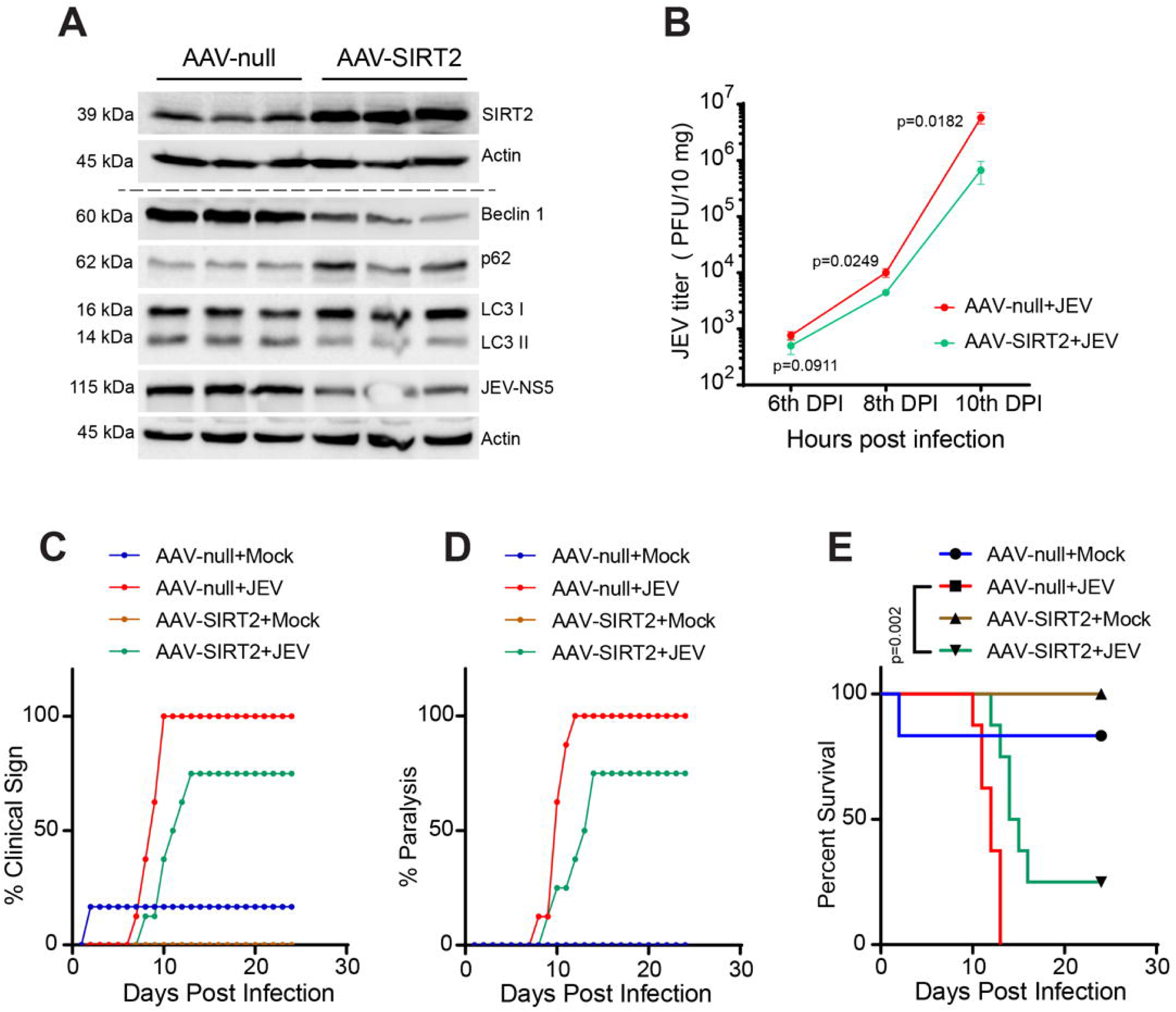
SIRT2 gene therapy using the AAV vector protects against JEV infection in mice. (**A-E**) 10-day-old wild-type mice were inoculated with 10^5^ PFUs of JEV through the foot-pad route, and 10^15^ copies of AAV-null or AAV-SIRT2 treatment started from first dpi and 4^th^ dpi through the IP route. (**A**) Representative immune-blot displaying changes in SIRT2 protein and autophagy markers in the brain of AAV-null or AAV-SIRT2 treated wild-type mice infected with JEV at 8^th^ dpi (n=3 mice each group). (**B**) The graph represents JE virus PFUs in the brain of JEV-infected wild-type mice, with AAV-null or AAV-SIRT2 treatment; n=3 mice per dpi, respectively. The p-value was calculated using Student’s t-test. The graph represents the percentage of animals exhibiting (**C**) clinical signs and (**D**) paralysis in mock or JE virus-infected wild-type mice treated with AAV-null or AAV-SIRT2 at various dpi (n=6-8 mice in each group). (**E**) Kaplan-Meier survival curve indicating the survival percentage of mock or JE virus-infected wild-type mice, with AAV-null or AAV-SIRT2 treatment at various dpi up to 24^th^ dpi (n=6-8 mice in each group). The p-value was calculated using the Log-rank (Mantel-Cox) test.

### SIRT2 deficiency regulates NF-κB transcriptional factor activity to enhance the inflammatory response upon JEV infection in mice brains

To understand the pathways deregulated upon JEV infection, we performed global mRNA-sequencing of mock and virus-infected WT mice brains. Our global transcriptome analysis demonstrated that JEV infection led to the deregulation of several genes including those involved in immune response (**Figure 5A; Figure S2A**). We also analyzed the transcription factors associated with the differentially expressed genes (**Figure 5B; Figure S2B**). According to our KEGG pathway analysis, cytokine and its receptor interaction was the most significantly upregulated pathway (**Figure 5C**). The transcription factor corresponding to the most significantly upregulated genes was NF-κB (**Figure 5D**). We also identified downregulated pathways and the associated transcription factors (**Figure 5E-5F**). Further, we analyzed transcriptomic data to determine the expression levels of approximately 400 genes regulated by NF-κB transcription factor^54^. Our analysis revealed that several NF-κB target cytokines and chemokines were transcriptionally upregulated in JEV-infected mice brains (**Figure 5G-5H; Figure S3A, S3B**). Further, a comparison of transcriptomic data between JEV-infected WT and SIRT2-KO mice brains revealed a significant increase in several NF-κB target inflammatory cytokines and chemokine genes (**Figure 5I-5J; Figure S4A, S4B**). Next, we validated the global transcriptomics results using quantitative PCR. We observed increased transcript levels of NF-κB target inflammatory cytokines and chemokine genes in JEV-infected mice brains. These transcripts were further increased in the JEV-infected SIRT2-KO mice brains (**Figure 5K-5N; Figure S5A-S5F**). During JEV infection, neuronal damage occurs (i) directly by virus replication in neurons and (ii) directly by JEV-induced increased cytokines in the brain^55–57^. Therefore, the increased susceptibility of SIRT2-KO mice to JEV infection could be attributed to both cytokine release due to hyperactivation of NF-κB and increased replication of the JEV in the brain.

**Figure 5.**
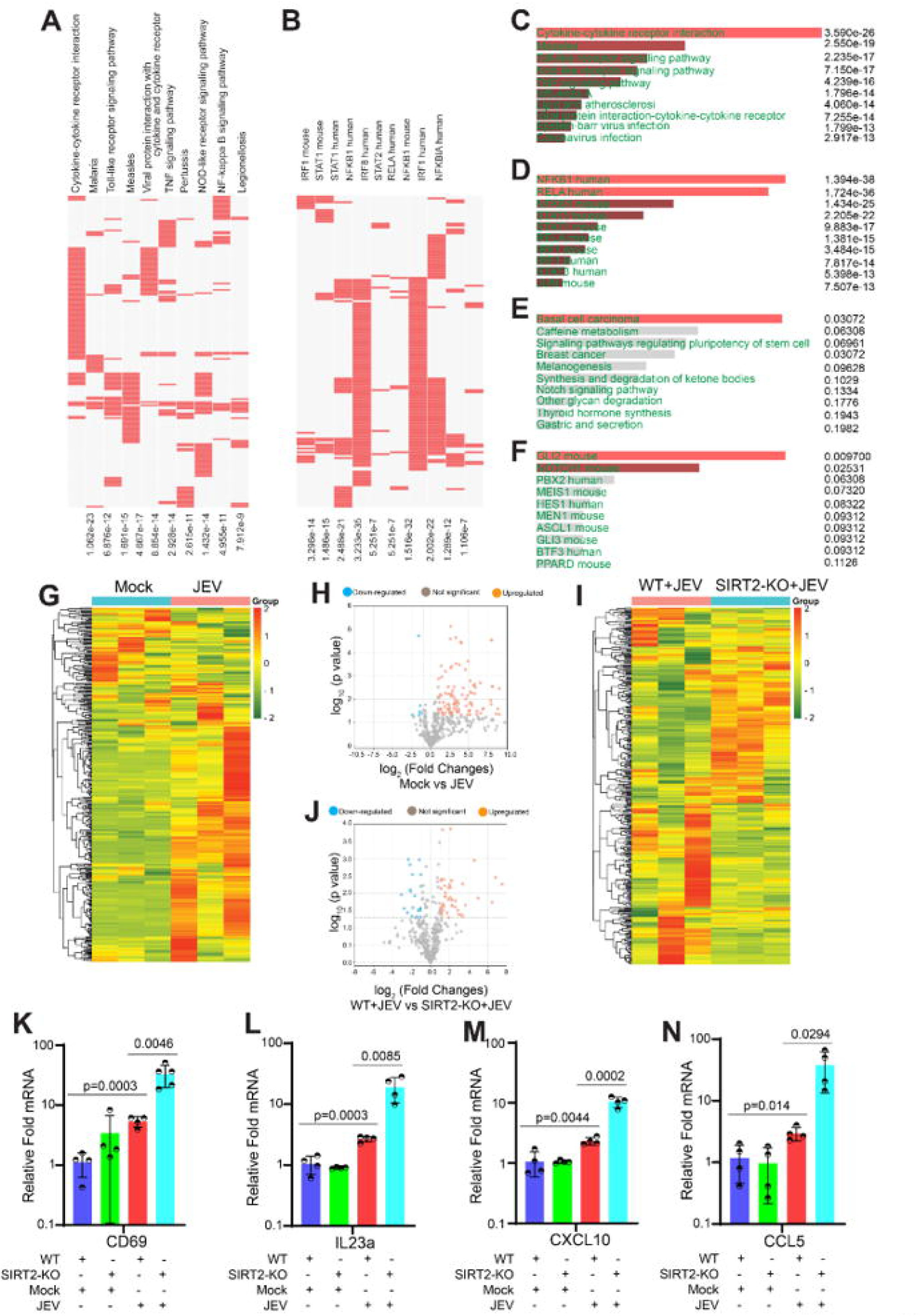
Hyperactivation of NF-κB transcriptional activity-dependent genes in SIRT2 knock-out mice upon JEV infection. (A-J) Transcriptome was performed in the mice brains infected with JE virus of 10^5^ PFUs by the foot-pad route. n=3 mice each group. (**A**) Clustergram representing the top hit KEGG pathway enrichment based on the genes significantly differentially expressed in the mRNA transcriptome of mock or JE virus infected WT mice brain (The differential expression gene names associated with the top hit KEGG pathway enrichment were illustrated in the enlarged figure in Figure S2 for better clarity).(**B**) Clustergram representing the top hit TRRUST Transcription Factors enrichment based on the genes significantly differentially expressed in the mRNA transcriptome of mock or JE virus infected WT mice brain (The d differential expression gene names associated with the top hit TRRUST Transcription Factors enrichment were illustrated in the enlarged figure in Figure S2 for better clarity). (**C**) Graph depicting top hit KEGG pathways enrichment based on the genes significantly upregulated in the transcriptome of JE virus infected mice brain compared to mock-infected WT mice. (**D**) Graph depicting top hit TRRUST Transcription Factors enrichment based on the genes significantly upregulated in the transcriptome of JE virus infected mice brain compared to mock-infected WT mice. (**E**) Graph showing top hit KEGG pathways enrichment based on the genes significantly down-regulated in the transcriptome of JE virus infected mice brain compared to mock-infected WT mice. (**F**) Graph depicting top hit TRRUST Transcription Factors enrichment based on the genes significantly down-regulated in the transcriptome of JE virus infected mice brain compared to mock-infected WT mice. (**G**) Heatmap representing differential expression profile of genes regulated by NF-κB in the transcriptome of mock or JE virus infected WT mice brain (The differential expression gene names associated with the NF-κB transcription factor activity were illustrated in the enlarged figure in Figure S3 for better clarity). (**H**) Volcano plot displaying differentially expressed profile of genes regulated by NF-κB transcriptional activity in the transcriptome of mock or JE virus-infected WT mice brain with log2 fold change ≥2 and p ≤0.05(n=3 mice each group) (The differential expression gene names associated with the NF-κB transcription factor activity were illustrated in the enlarged figure in Figure S3 for better clarity). (**I**) Heatmap representing differential expression profile of genes regulated by NF-κB transcriptional activity in the transcriptome of JE virus-infected WT or SIRT2^-/-^ mice brain (The differential expression gene names associated with the NF-κB transcription factor activity were illustrated in the enlarged figure in Figure S4 for better clarity). (**J**) Volcano plot displaying differential expression profile of genes regulated by NF-κB transcriptional activity in the transcriptome of JE virus-infected WT or SIRT2^-/-^ mice brain with log2 fold change ≥2 and p ≤0.05 (The differential expression gene names associated with the NF-κB transcription factor activity were illustrated in the enlarged figure in Figure S4 for better clarity). (**K-N**) Ten-day-old mice pups were infected through the foot-pad route with 10^5^ PFUs of JE virus. Relative fold change in the (**K**) CD69; (**L**) IL23a; (**M**) CXCL10; and (**N**) CCL5 mRNA levels on 8^th^ dpi of mock or JE virus-infected WT or SIRT2^-/-^ mice brains; n=4 mice per group; P value calculated using Student’s t-test.

### SIRT2 mediated acetylation of NF-κB regulates JEV replication through NF-κB-autophagy axis

Since our findings indicate that SIRT2 regulates JEV replication in neuro2a cells and mice brains (**Figure 2A-2B; Figure S1A-S1B; Figure 2F**; **Figure 2K**; **Figure 3A**; **Figure 3F**) with an increase in NF-κB target gene expression in JEV-infected SIRT2-KO mice brain (**Figure 5A-5N; Figure S5A-S5C**), we checked if SIRT2 directly interacts with the NF-κB and regulate its activity upon JEV infection. To investigate SIRT2-NF-κB protein-protein interaction, we immunoprecipitated NF-κB from mock or JEV-infected neuro2a cells. We identified SIRT2 interacts with NF-κB, and the interaction is reduced upon JEV infection (**Figure 6A**). Further, JEV infection increased acetylation levels of NF-κB (**Figure 6A**). Furthermore, we immunoprecipitated NF-κB from JEV-infected neuro2a cells treated with Ad-null or Ad-SIRT2 and observed reduced NF-κB acetylation and JEV-NS5 levels in the SIRT2-overexpressing cells (**Figure S6**). Next, we performed a reporter assay to evaluate the transcriptional activity of NF-κB. In line with transcriptome analysis, we found that JEV infection increased NF-κB transcriptional activity (**Figure 6B**). Interestingly, SIRT2 deficiency enhanced JEV-infection mediated NF-κB transcriptional activity (**Figure 6C**) and increased the viral load in neuro2a cells (**Figure 2B**). Conversely, SIRT2 overexpression significantly reduced the JEV-infection-induced increase in NF-κB transcriptional activity (**Figure 6D**). It also reduced the viral load in neuro2a cells (**Figure S1A, S1B**). Taken together, these findings suggest that JEV infection negatively regulates SIRT2 levels, resulting in a concomitant increase in acetylated NF-κB mediated virus replication, and SIRT2 overexpression reverts this phenotype. Next, we attempted to identify the cellular signaling pathways related to NF-κB target genes that may be involved in JEV replication. We compared the transcriptome profiles of JEV-infected WT and SIRT2-KO mice brains and carried out gene network analysis. We observed that NF-κB mediated inflammation and autophagy were among the key axes that were deregulated in JEV-infected SIRT2-KO brains compared to JEV-infected WT mice brains (**Figure 6E**). Notably, NF-κB is established to induce autophagy by transcriptional upregulation of beclin-1^58^. Our previous study has identified a crucial role for autophagy in JEV replication^1^. Therefore, we explored the possibility of the SIRT2-NF-κB-beclin-1 axis of autophagy in JEV infection. We observed that JEV infection significantly upregulated the transcript levels of beclin-1 (**Figure 6F**), in concordance with the increased NF-κB transcriptional activity (**Figure 6B**). Additionally, SIRT2-KD also enhanced the transcript levels of beclin-1 (**Figure 6G**) and NF-κB transcriptional activity (**Figure 6C**). Conversely, SIRT2-OE significantly reduced the transcript levels of beclin-1 (**Figure 6H**) and NF-κB transcriptional activity (**Figure 6D**) and caused a reduction of viral load in neuro2a cells (**Figure S1A-S1B**). Further, we found that neuro2a cells infected with JEV showed increased autophagy as indicated by autophagy markers such as increased beclin-1, increased LC3II and reduced p62 (**Figure 6I; Figure S7A**). In line with our *in vitro* findings, JEV-infected mice brains also showed increased autophagy through increased beclin-1, LC3II, and reduced p62 concomitant with reduced SIRT2 levels (**Figure 6J; Figure S7B**). Further, we found that SIRT2 knockdown as well as chemical inhibition using AGK2 increased beclin-1 and autophagy in JEV-infected neuro2a cells (**Figure 6K; Figure S7C**; **Figure 6L; Figure S7D**) and mice brains (**Figure 3F-3G**). Moreover, JEV-infected SIRT2-KO mice showed increased beclin-1 and autophagy compared to their WT counterparts (**Figure 2E**; **Figure 6M; Figure S7E**). In contrast, SIRT2 overexpression significantly reduced autophagy and inhibited viral multiplication in neuro2a cells (**Figure S1A, S1B; Figure 6N; Figure S7F**). Moreover, we observed that AAV-mediated SIRT2 gene therapy down-regulated the JEV-induced autophagy, which is critical for JEV replication, thereby exhibiting an antiviral role in the JEV-infected mice (**Figure 4A**). These results suggest that SIRT2 regulates JEV replication through the NF-κB-autophagy axis.

**Figure 6.**
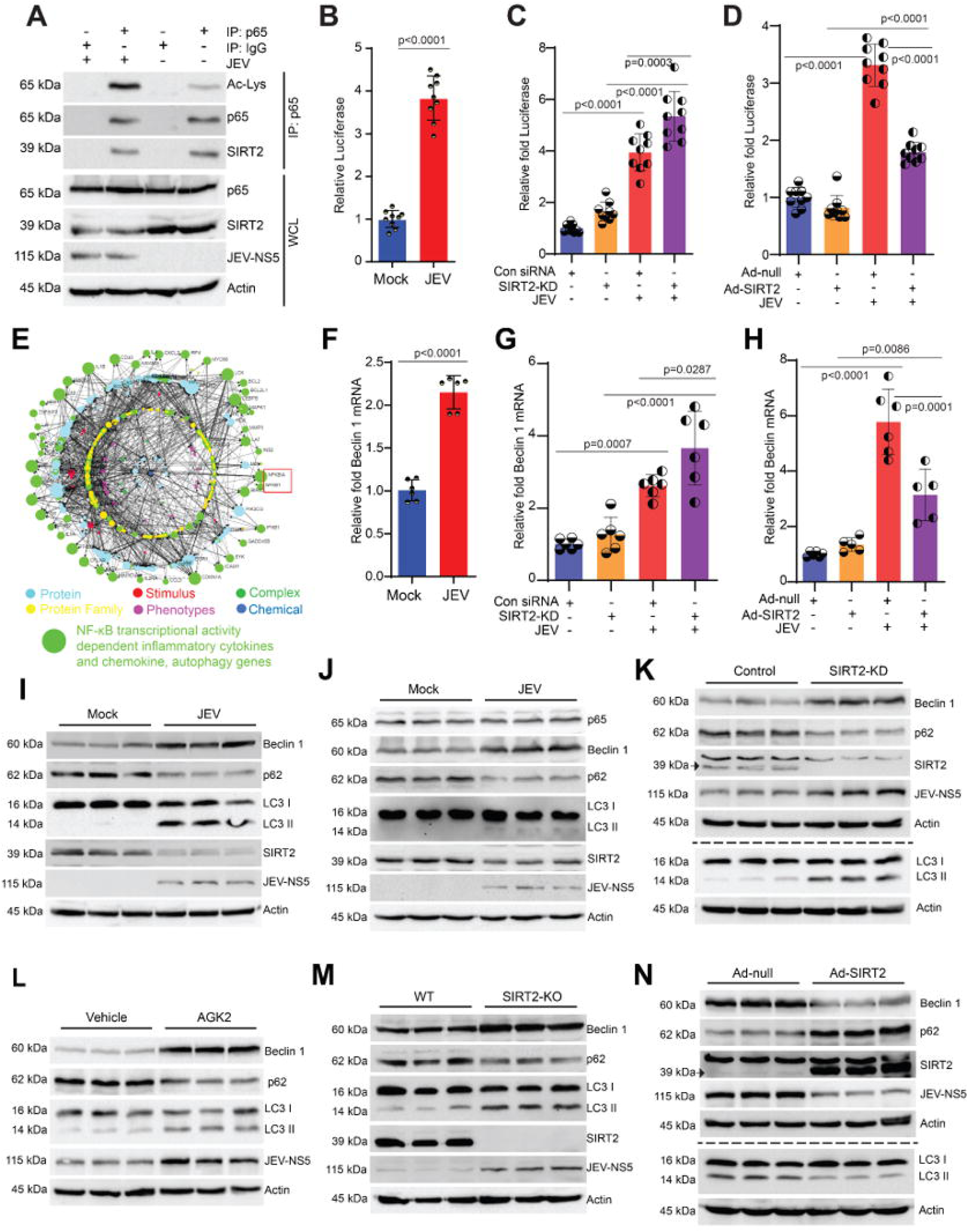
SIRT2 and NF-κB interaction mediated acetylation status of NF-κB regulates the JEV replication. (**A**) Immunoprecipitation of p65 at 30 hpi with mock or JE virus (1 MOI) infected neuro2a cells demonstrating p65 and SIRT2 interaction. (**B**) NF-κB activity was assayed at 30 hpi using luciferase reporter in mock or JE virus (1 MOI) infected neuro2a cells (n=3 independent experiments, triplicates were used in each experiment; The p-value was calculated using Student’s t-test). Relative fold change in the luciferase reporter activity of NF-κB at 30 hpi of JE virus infection in (**C**) control or SIRT2-KD; (**D**) Ad-null or Ad-SIRT2; treated conditions; n=3 independent experiments, triplicates were used in each experiment; Statistical analysis using one-way ANOVA. **(E)** The network of signaling pathways represents differential expression genes that are regulated by NF-κB transcriptional activity in the WT or SIRT2^-/-^ mice brain upon JEV infection (n=3 mice each group). The qPCR analysis representing the relative fold changes in the beclin-1 expression levels in (**F**) JE virus-infected (1 MOI) neuro2a cells; (**G**) JEV (1 MOI) infected control or SIRT2-KD neuro2a cells; and (**H**) JEV (1 MOI) infected Ad-null or Ad-SIRT2 treated neuro2a cells at 30 hpi (n=3 independent experiments, triplicates were used in each experiment; The p-value was calculated using (B and F) Student’s t-test and (C-D and G-H) one-way ANOVA. **(I)** Representative immuno-blotting images displaying differences in autophagy markers at 30 hpi in the JE virus-infected neuro2a cells. (n=3 independent experiments; duplicates were used in each experiment). **(J)** Representative immuno-blotting images displaying an increase in autophagy markers at 8 dpi in the JE virus-infected WT mice brain (n=6 mice in each group). **(K)** Representative immuno-blotting images displaying differences in autophagy markers at 30 hpi in the JE virus-infected, control or SIRT2-KD neuro2a cells (n=3 independent experiments; duplicates were used in each experiment). **(L)** Representative immuno-blotting images displaying change in autophagy markers at 30 hpi in the JE virus-infected, vehicle or AGK2-treated neuro2a cells (n=3 independent experiments; duplicates were used in each experiment). **(M)** Representative immuno-blotting images displaying differences in autophagy markers at 8 dpi in the WT or SIRT2^-/-^ mice brains upon JEV infection (n=6 mice in each group). **(N)** Representative immuno-blotting images displaying changes in autophagy markers at 30 hpi in the JE virus-infected neuro2a cells upon Ad-null or Ad-SIRT2 treatment (n=3 independent experiments; duplicates were used in each experiment). Statistical analysis of all immune-blotting experiments was performed using Student’s t-test.

### SIRT2 deficiency promotes JEV replication in mice through NF-κB activity

Previous studies indicate that NF-κB activity is negatively regulated by NF-κB inhibitor, IκB, and this negative regulation is released by phosphorylation of IκB^50,51^. Thus, we treated the JEV-infected neuro2a cells with Bay-11, an inhibitor of IκBα phosphorylation, and measured NF-κB activity using a luciferase activity-based reporter assay. We observed a significant reduction in JEV-induced NF-κB activity (**Figure 7A**) and a reduction in the viral load in neuro2a cells upon Bay-11 treatment (**Figure 7B**), indicating that IκBα phosphorylation-mediated activation of NF-κB activity is required for JEV replication. Moreover, Bay-11 further decreased NF-κB activity and JEV load in SIRT2 depleted neuro2a cells as compared to controls (**Figure 7C, 7D**). In contrast, SIRT2 overexpression remarkably reduced JEV-induced NF-κB transcription factor activity and JEV load which were reduced further upon Bay-11 treatment (**Figure 7E, 7F**). Interestingly, Bay-11 significantly reduced SIRT2-KD mediated increase in NF-κB transcriptional activity under JEV-infected conditions (**Figure 7C**). Similarly, Bay-11 significantly reduced viral load which was observed upon SIRT2-KD in neuro2a cells (**Figure 7D**). Overall, these results reveal that SIRT2 exerts an anti-viral effect by deacetylating and inhibiting NF-κB activity, which is required for JEV replication.

**Figure 7.**
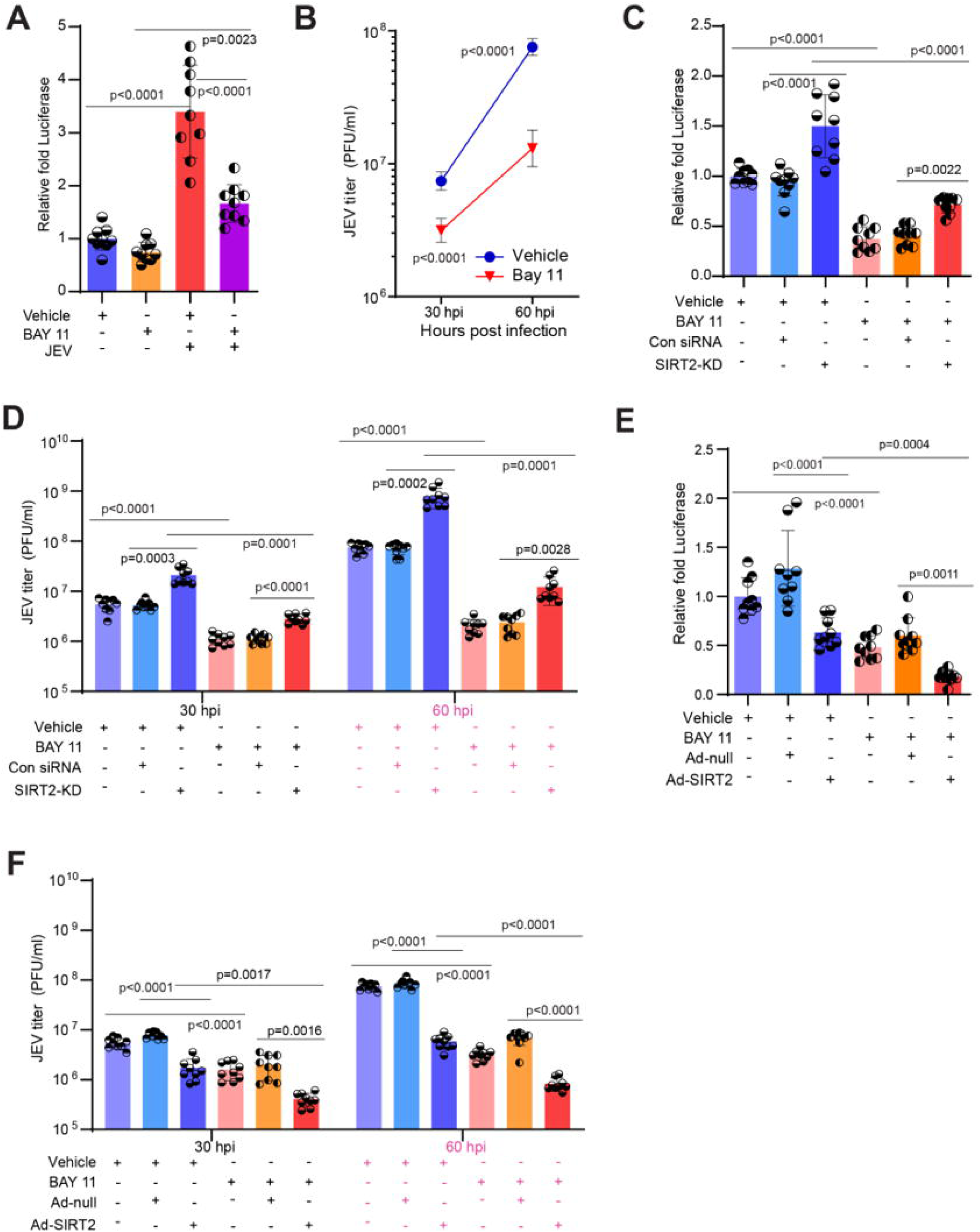
SIRT2 exerts the antiviral effect by inhibiting NF-κB transcriptional activity. **(A**) Graph displaying NF-κB activity assessed at 30 hpi using luciferase reporter assay in mock or JE virus (1 MOI) infected neuro2a cells treated with vehicle or Bay-11. Results are expressed as fold change relative to the vehicle-treated mock control group. (n=3 independent experiments, triplicates were used in each experiment; p-values are calculated using one-way ANOVA. (**B**) Graph corresponding to changes in viral PFUs at 30 and 60 hpi in JE virus (1 MOI) infected neuro2a cells with vehicle or Bay-11 treatment. n=3 independent experiments, triplicates were used in each experiment; The p-value was calculated using Student’s t-test. **(C)** Graphical representations of NF-κB activity evaluated by luciferase reporter assays in JE virus (1 MOI) infected neuro2a cells treated with vehicle or Bay-11 in Control siRNA or SIRT2 specific siRNA transfected neuro2a cells; Results are depicted as fold change relative to vehicle-treated non-transfected control groups. n=3 independent experiments, triplicates were used in each experiment; Statistical analysis using one-way ANOVA. (**D**) Graph displaying changes in viral PFUs at 30 hpi in JEV infected neuro2a cells upon transfection with Control siRNA or SIRT2 specific siRNA and further treatment with vehicle or BAY-11. (n=3 independent experiments, triplicates were used in each experiment); p-values are calculated using one-way ANOVA. **(E)** Graphical representations of NF-κB activity evaluated by luciferase reporter assays in JE virus (1 MOI) infected neuro2a cells treated with vehicle or Bay-11 in Ad-null or Ad-SIRT2 treated neuro2a cells; Results are depicted as fold change relative to vehicle-treated non-transfected control groups. n=3 independent experiments, triplicates were used in each experiment; Statistical analysis using one-way ANOVA. **(F)** Graph displaying changes in viral PFUs at 30 hpi in JEV infected neuro2a cells upon treatment with Ad-null or Ad-SIRT2 and further treatment with vehicle or BAY-11. (n=3 independent experiments, triplicates were used in each experiment); p-values are calculated using one-way ANOVA.

Subsequently, to determine whether SIRT2 regulates JEV replication in mice brains through NF-κB, JEV-infected (through foot pad route of infection) WT and SIRT2-KO mice were treated with BAY-11. Here, we observed that BAY-11 treatment in JEV-infected WT mice reduced the viral load in both serum and brain, thereby reducing the clinical outcomes including paralysis, and enhancing survival compared to vehicle-treated JEV-infected WT mice (**Figure 8A-8E**). Furthermore, to study the therapeutic potential of BAY-11, we administered BAY-11 to the JEV-infected mice after the onset of JEV clinical signs. Even when administered after the onset of JEV clinical signs in mice, BAY-11 significantly reduced paralysis (**Figure S8A**), improved the survival percentage of mice infected with JEV (**Figure S8B**), and reduced the JEV load in the mice brains (**Figure S8C**). These results indicate that IκBα phosphorylation mediated activation of NF-κB is required for JEV replication in mice brains. Besides this, BAY-11 treatment in JEV-infected SIRT2-KO mice rescues the exacerbated clinical signs, paralysis, and decline in survival rates observed in the JEV-infected SIRT2-KO mice (**Figure 8A-8C**). Additionally, BAY-11 treatment reduced the increased viral load observed in JEV-infected SIRT2-KO mice serum and brain (**Figure 8D, 8E**). Collectively, our results suggest that SIRT2 deficiency in mice exacerbates JEV induced clinical signs, paralysis, and causes decline in survival rates through the activation of the NF-κB transcription factor.

**Figure 8.**
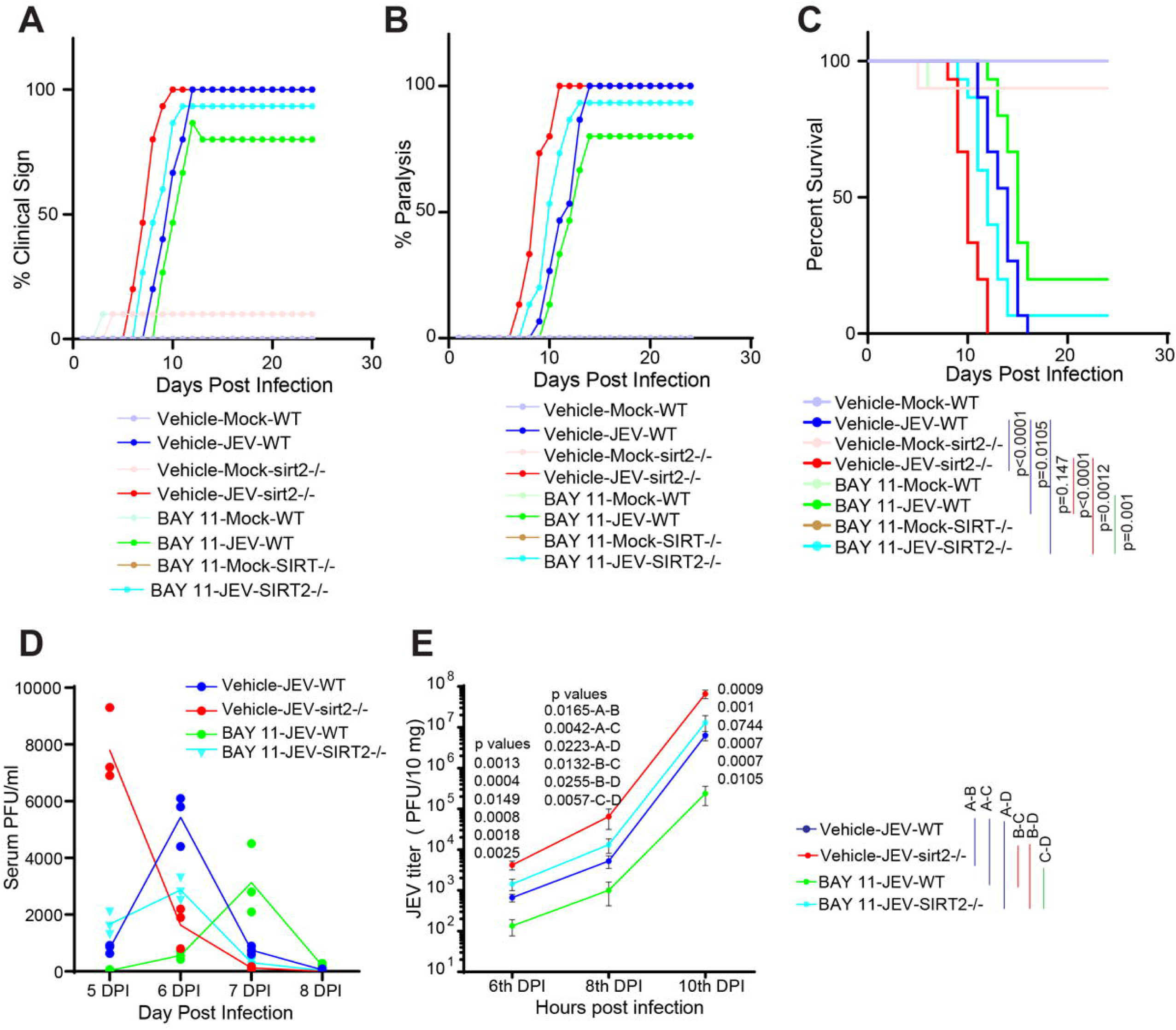
SIRT2 regulates the NF-κB mediated antiviral activity in JE virus-infected mice. (**A-E**) 10-day-old WT or SIRT2^-/-^ mice were inoculated with 10^5^ PFUs of JEV through the foot-pad route, and vehicle or BAY-11 treatment started from first dpi through the IP route for every day up to 24^th^ dpi. The graph represents the percentage of WT or SIRT2^-/-^ animals exhibiting (**A**) clinical signs (**B**) paralysis in JE virus-infected vehicle or BAY-11 treated mice at various dpi (n=10-15 mice in each group). (**C**) Kaplan-Meier survival curve indicating the survival percentage of JE virus-infected, WT or SIRT2^-/-^ mice, with vehicle or BAY-11 treatment at various dpi up to 24^th^ dpi (n=10-15 mice each group). The p-value was calculated using the Log-rank (Mantel-Cox) test. The graph represents JE virus PFUs in the (**D**) serum and (**E**) brain of infected WT or SIRT2^-/-^ mice, with vehicle or BAY-11 treatment; n=3 and 5 mice per dpi, respectively. The p-value was calculated using Student’s t-test.

## Discussion

While vaccines play an important role in preventing virus infection, antivirals are needed to complement the limitations of vaccines. Identifying host-directed antivirals is important as they reduce the resistance acquired by virus mutations^59–62^. This present study describes SIRT2 as a host-directed antiviral factor against JEV, a public health threat due to the lack of specific complement antivirals in clinical practice despite the availability of vaccines. Notably, the existing studies have shown that SIRT2 knockdown increases the replication of a few DNA and RNA viruses in cell culture^14^. The present study elucidates that SIRT2 genetic deficiency or SIRT2 chemical inhibitors enhance JEV replication in cell culture and mouse models, while SIRT2 overexpression displays antiviral characteristics. Moreover, the present study established that SIRT2 acts as a host-directed antiviral by inhibiting JEV-induced autophagy by regulating the NF-κB-Beclin-1-autophagy axis, thereby reducing virus replication.

SIRT2 is established to transcriptionally regulate several cellular signalling pathways through deacetylating transcriptional factors^58,63–69^. In the case of DNA viruses that replicate in the nucleus, SIRT2 regulates transcriptional factors p53^70^, and deacetylates G3BP, thereby regulating cGAS STING-IRF3 transcriptional activity^19^. Besides this, SIRT2 was reported to transcriptionally regulate autophagy by regulating transcriptional factors such as FoxO1 and TFEB^71,72^. The present study reveals that NF-κB deacetylation through SIRT2-NF-κB interaction inhibits virus-induced autophagy. Mechanistically, SIRT2-mediated deacetylation of NF-κB inhibits the transcriptional activity of the NF-κB, thereby inhibiting the transcription of Beclin-1 and subsequent downstream autophagy, which is crucial for virus replication. The NF-κB transcriptional activity regulates approximately 400 genes^58,68,73^. On the other hand, Beclin-1 is reported to maintain the persistent activity of NF-κB in human T cell leukaemia virus type 1 transformed T lymphocytes^74^. Previous studies have reported that Beclin-1 regulates flavivirus-induced autophagy by viral non-structural protein 1 (NS1)^75^ and mosquito saliva-specific protein allergen-1^76^, indicating that Beclin-1 is needed for autophagy induction to favour flavivirus replication. Along these lines, our previous study has found that JEV infection enhances virus replication by inducing autophagy through the transcriptional factor PTEN^1^. Our findings unveil that JEV infection downregulates SIRT2, which causes increased acetylation of NF-κB, leading to the activation of the NF-κB pathway. Furthermore, we observed that the activated NF-κB pathway by JEV infection induces autophagy through Beclin-1, thereby enhancing virus replication. Previous studies suggest both pro- and anti-inflammatory roles for SIRT2 in different pathologies. SIRT2 inhibition with AGK2 conferred protection against NF-kB and MAPK activation mediated inflammatory response in the liver of mice model of thioacetamide-induced acute liver failure^77^. Similarly, in other studies, including the lethal septic model and LPS-induced renal inflammation, SIRT2 inhibition attenuated inflammation^78,79^. On the other hand, the anti-inflammatory role of SIRT2 is reported in traumatic brain injury and neuropathic pain^80,81^. However, our findings show that SIRT2 depletion or chemical inhibition exacerbates inflammatory response and prognosis in mice models of JEV infection whereas SIRT2 gene therapy confers protection to mice against JEV infection. Previous findings suggest that the severity of JEV infection is largely manifested by inflammatory cytokine and chemokine hyperactivation^57,82–84^. In the current study, we observed that the exacerbated clinical outcome in JEV-infected SIRT2-KO mice was mediated by NF-κB transcriptional activity-dependent hyperactivation of inflammatory cytokines and chemokines.

Our findings establish that SIRT2 negatively regulates NF-κB transcriptional activity and exhibits antiviral activity by restricting virus replication and reducing disease severity. We believe that SIRT2 gene therapy could be taken forward to the next level of clinical studies, which may improve the treatment of JEV infection. Our findings stress the potential therapeutic implications for RNA viral infections that utilize the evolutionarily conserved mechanism of the autophagy axis for favouring virus replication and inflammatory cytokines and chemokines hyperactivation for pathogenesis.

## Materials and Methods

### Experimental animals

In the present study, the SIRT2^+/+^ and SIRT2^-/-^ mice were procured from the Jackson Laboratory (B6.129-Sirt2tm1.1Fwa/J, RRID: IMSR_JAX:012772). The mice were housed and bred in the institute’s central animal facility at Indian Institute of Science, Bengaluru, India. The mice were maintained in individually ventilated cages, with a 12-hour light/dark cycle. Food and water were provided ad libitum. All experiments were conducted after approvals from the institutional animal ethics committee and the guidelines of the Committee for the Purpose of Control and Supervision of Experiments on Animals, Government of India. Further, all the studies followed as per the U. S. National Institutes of Health (NIH Publication 2011) published 8^th^ edition of Care and Use of Laboratory Animals.

### JEV viral strain

Japanese encephalitis virus (JEV, Vellore strain-P20778) used in the study was a kind gift from Professor P. N. Rangarajan, Division of Biological Science, Department of Biochemistry, Indian Institute of Science (IISc), Bengaluru, India.

### Method details

#### Mice model for JEV infection

The 10-day-old mice were infected with JEV by two routes, including foot-pad route and intra-cerebral infection at 10^5^ and 3x10^3^ PFU, respectively, and continuously observed and scored for the clinical signs, paralysis, and survival every day for 24 days of post-infection. Inhibitors AGK2 and BAY-11 were administered intraperitoneally at 2.5 and 2.5 mg/kg/day, respectively.

#### Cell culture, Cell transfection, and drug treatment

The *in vitro* studies were performed on neuro2a cells. The cells were cultured in high glucose DMEM supplemented with 10% foetal bovine serum (FBS) and an antimycotic-antibiotic mix and grown and maintained in a humidified incubator at 37°C and 5% CO_2_. The siRNA-based SIRT2 knockdown and adenovirus-based overexpression of SIRT2 was carried out at 70-80% confluency as described previously^67,85,86^. Briefly, siRNA transfection was carried out using Lipofectamine RNAiMAX per the manufacturer’s protocol. In the case of adenovirus transduction, SIRT2 or null adenovirus, propagated and quantified in the HEK 293T cell line, was used with equal MOIs to infect the neuro2a cells. AGK2 and BAY 11-7082 were used to inhibit SIRT2 and IKK inhibitors at 10 µM and 2.5 µM concentrations, respectively.

#### JEV propagation and viral plaque assay

The BHK-21 cell line was used to propagate JEV and measure the plaque-forming units (PFU) as described previously^1^. At a density of 0.2x10 BHK-21 cells per well, cells were seeded in six-well plates, and infected with ten-fold serial dilutions of JEV virus for 1 hour for the virus entry. After an hour, the uninfected viruses were removed by washing wells thrice using phosphate-buffered saline (PBS). Then cells were covered using 3% low-melting agarose in culture DMEM with 10% FBS and antibiotic-antimycotic. Further, it was allowed at room temperature for 15 minutes to solidify. After solidification, incubated in a humidified incubator at 37°C and 5% of CO_2_ for 36-48 hours. Then the cells were examined microscopically after fixing using 10% formaldehyde for 10-20 minutes at room temperature and stained with a solution of 1.5% crystal violet. Then, the excess strains were removed by washing thrice with MilliQ water, and the plaques were counted using white light. According to the plague counts and virus serial dilution used, PFU was calculated per mL as follows: PFU=N x DF/V (DF is virus dilution factor, N is number of plaques, and V is the volume of the inoculum.

#### Dual-Luciferase reporter assay

To study the NF-κB activity, the cells (neuro2a) were co-transfected with plasmids containing NF-κB-Luciferase reporter and Renilla-Luciferase reporter with pRL-CMV using lipofectamine 3000. While NF-κB-Luciferase reporter activity corresponded to NF-κB activity, Renilla-Luciferase activity was used to normalise transfection. The cell lysis and assay were done in accordance with the Dual-Luciferase Reporter kit Promega, and Pharmingen Moonlight 3010, BD Biosciences, was used to measure the luminescence.

#### Cyto-immunofluorescence microscopy

Cyto-immunofluorescence microscopy was carried out as described previously^1^. In brief, the cells were cultured on the coverslips and subjected to mentioned treatments. At the study time point, the cells were fixed by incubating with 10% formaldehyde for about 15-20 minutes at room temperature (RT). The fixed cells were washed thrice with PBS and then permeabilization was carried out using 0.25% Triton X-100 diluted in PBS by incubating at RT for 10-15 min. The permeabilized cells were blocked for 20 to 30 minutes with 1% BSA dissolved in PBS with 0.1% tween and 22.52 mg/mL glycine. The blocked cells were then incubated with diluted primary antibody at 4°C overnight. Post-incubation, the cells were washed thrice and incubated with diluted secondary antibodies conjugated with Alexa Fluor for an hour at RT. The nuclear stain Hoechst 33342 was also incubated with a secondary antibody. After secondary incubation, the coverslips were washed and mounted on a glass slide with a Fluoromount-G Aqueous mount. The cells were imaged using a confocal microscope Zeiss LSM 880.

#### Immunoprecipitation assay

Immunoprecipitation (IP) subsequently immunoblotting was done as described previously^1^. In brief, the neuro2a cells, after aspiration of media, were washed thrice with PBS and cell lysates were prepared in RIPA cell lysis buffer. The protein levels were quantified using Bradford, and about 500 μg protein was used for IP. The lysate containing the specified protein levels was incubated with 2 μg of specific or control (IgG)) antibodies rotating at 5 rpm overnight. After overnight incubation, washed protein A/G magnetic beads were added to the lysate containing antibodies and incubated at 4°C for 2 hours. Post-incubation, the entire mixture was passed through the column, washed thrice using PBS, and finally, elution was carried out using Laemmli buffer. The IP was measured by immunoblotting.

#### Real-time PCR

For the real-time PCR, TRI-reagent was used to extract RNA from the cells/tissues according to the manufacturer’s guidelines. Then, the extracted RNAs were quantified in nanodrop, and 500 ng was used for cDNA synthesis utilizing the iScript cDNA Synthesis Kit. The real-time PCR was carried out using Quant studio 6 flex using the qPCR master mix with SYBR-Green.

#### Immuno-blotting

Immunoblotting was performed as described previously^1^. In Brief, the cell/tissue lysates were prepared using RIPA buffer with phosphatase inhibitor, deacetylase inhibitors and protease inhibitor cocktail. The tissue samples were homogenized with liquid nitrogen and then collected in RIPA buffer. The lysates were centrifuged for 15 minutes at 12,000 rpm. The supernatants were collected separately and used for western blotting. The protein quantities were measured by using a Bradford reagent. An equal amount of protein for each sample was loaded on 12% SDS-PAGE and resolved followed by transfer to a PVDF membrane (0. 45μm) by overnight transfer method. Upon completion of the transfer, the PVDF membrane (0. 45μm) was incubated in 5% skimmed milk in Tris-buffered saline with Tween-20 (TBST) for an hour at RT. Then, the PVDF membrane was washed using TBST for thrice, three minutes each and incubated with the primary antibody of the mentioned dilutions at 4°C for overnight. Post-incubation, the unbound antibodies were washed off with three TBST washes for three minutes each. The membrane was incubated with appropriate secondary antibodies conjugated to horseradish peroxide for an hour at RT. The chemiluminescence was detected with ECL reagents, and the image was attained in a chemiluminescence imager.

#### Generation of adeno-associated virus

Mouse full-length SIRT2 cDNA was cloned into the AAV-CMV-GFP (Plasmid #67634)^87^ by replacing GFP to generate AAV-CMV-mSIRT2. The pAAV2/9n (Plasmid #112865)^88^ was used RepCap plasmids by inserting WVLPSGG between amino acids 588 and 589, corresponding to AAV9 VP1^89^ in this plasmid by using the QuikChange Site-Directed Mutagenesis Kit^90^. The pAdDeltaF6 (Plasmid #112867)^91^ was used as an AAV helper plasmid. The AAV-CMV-mSIRT2, pAdDeltaF6, and WVLPSGG-inserted pAAV2/9n were co-transfected in the HEK 293T cell line to generate the AAV vector to deliver SIRT2 into mice’s brains. The control null-AAV vector was generated by deleting the SIRT2 in the AAV-CMV-mSIRT2 and co-transfected with pAdDeltaF6, and WVLPSGG-inserted pAAV2/9n in the HEK 293T cell line. The resulting AAV vector was purified by ultracentrifugation as per the previous protocol^87,92,93^. The AAV genomic DNA copy number was determined by qPCR. Equal copies of purified SIRT2-AAV and null-AAV were used to infect the mice through the intraperitoneal route.

#### Global mRNA sequencing and analysis

Briefly, the total RNA was isolated for transcriptome sequencing using TRI-reagent from the mock or JEV-infected SIRT2^+/+^ and SIRT2^-/-^ mice brains. The quality control was done with the 4150 Tape Station-System, Agilent-G2992AA with RNA ScreenTape-System, Agilent, 5067-5576. Samples with the above 8 RNA integrity number (RINe) values were utilized to further subsequent downstream processing. To enrich for mRNA in the samples, NEBNext Poly (A) mRNA magnetic isolation module was used and library preparation was carried out using 500ng of total RNA as per the manufacturer’s protocol, New England Biolabs-E7490 was employed. After the library preparation, the global mRNA sequencing was carried out in an Illumina-HiSeq sequencer. The obtained sequences were validated in FastQC (http://www.bioinformatics.babraham.ac.uk/projects/fastqc/). Then, fastp v0.20^94^ was used to remove the adapter sequences and low-quality reads. Next, STAR v2^95^ with Mus musculus database was used to map the genes. The feature-counts^96^ was used for mapped read counts and gene expression values. The genes that showed consistent differential expression when normalised using edgeR^97^. The gene network analysis was performed using SIGnaling Network Open Resource with SIGNOR 2.0. The KEGG pathways enrichment and TRRUST Transcription Factors enrichment analysis were carried out using Enrichr^98–100^.

### Statistical analysis

The statistical analysis and graphs included in the study were carried out in the GraphPad Prism 8.0, GraphPad Software Inc., USA. The mean±standard deviation was used to highlights in the graphs. The p-value was calculated using the student’s t-test, and one-way ANOVA, a <0.05 p-value was considered as statistically significant. For survival curve was determined using the Kaplan-Meier curve. The p values in the same were calculated using the Log-rank (Mantel-Cox) test. The ZEN image analysis software was used to analyse the confocal images. The western blots were quantified using ImageJ software.

### Data and code availability

The global mRNA-seq data from the mock or JEV-infected SIRT2^+/+^ and SIRT2^-/-^ mice brain has been submitted in the NCBI (PRJNA1001148 and PRJNA1091868). These data are accessible publicly from the date of manuscript publication.

## Acknowledgements

We are grateful to Prof. P.N. Rangarajan, Department of Biochemistry, IISc, Bangalore, India, for gifting the JE virus. The PAD received INSPIRE faculty (DST/INSPIRE/04/2016/001067) support for fellowship and research funds from the Department of Science and Technology, Government of India. P.A.D. received a research grant from the Science and Engineering Research Board (SERB) (CRG/2018/002192), Government of India.

## Author contributions

PAD and NRS conceived the idea of the study and planned all experimental conditions. PAD carried out almost all experiments. LD and VR assisted PAD in performing experiments. PAD, AKT and RSR were involved in the mice experiments. PAD and KM carried out the data arrangement. The first draft of the manuscript was written by PAD and KM. AS and SR engaged in data verification and manuscript review. The final version of the manuscript was written by PAD and NRS.

## Supplementary Figures and Legends

**Figure S1. SIRT2 overexpression reduces JEV viral titre in neuro2a cells.** (**A**) Representative western blotting confirming SIRT2 overexpression in neuro2a cells. **(B)** Graph displaying differences in JEV PFUs at the various hpi in JE virus-infected (one MOI) neuro2a cells upon treatment with Ad-null or ad-SIRT2. n=3 independent experiments; each experiment performed in triplicates; The p-value was calculated using Student’s t-test.

**Figure S2. Clustergram representing top hit KEGG pathways and TRRUST Transcription Factors enrichment.** (**A**) Clustergram representing the top hit KEGG pathways enrichment based on the genes significantly differentially expressed in the transcriptome of mock or JE virus-infected WT mice brain (n=3 mice each group). (**B**) Clustergram representing the top hit TRRUST Transcription Factors enrichment based on the genes significantly differentially expressed in the transcriptome of mock or JE virus-infected WT mice brain (n=3 mice each group).

**Figure S3. Heatmap and Volcano plot of NF-**κ**B transcriptional activity-dependent genes in the transcriptome of mock or JE virus-infected WT mice brain**. (**A**) Heatmap representing differential expression profile of genes regulated by NF-κB transcriptional activity in the transcriptome of mock or JE virus-infected WT mice brain (n=3 mice each group). (**B**) Volcano plot displaying differential expression profile of genes regulated by NF-κB transcriptional activity in the transcriptome of mock or JEV virus infected WT mice brain (n=3 mice each group) with log2 fold change ≥2 and p ≤0.05.

**Figure S4. Heatmap and Volcano plot of NF-**κ**B transcriptional activity-dependent genes in the transcriptome of JE virus-infected SIRT2^+/+^ or SIRT2^-/-^ mice brain.** (**A**) Heatmap representing differential expression profile of genes regulated by NF-κB transcriptional activity in the transcriptome of JE virus-infected SIRT2^+/+^ or SIRT2^-/-^ mice brain (n=3 mice each group). (**B**) Volcano plot displaying differential expression profile of genes regulated by NF-κB transcriptional activity in the transcriptome of JE virus-infected SIRT2^+/+^ or SIRT2^-/-^ mice brain (n=3 mice each group) with log2 fold change ≥2 and p ≤0.05.

**Figure S5. Relative fold change in the CD80, CXCL11, CLDN2, IFNb, IL1b, and IL18 mRNA levels in JE virus-infected SIRT2^+/+^ or SIRT2^-/-^ mice brain.** (**A-C**) 10-day-old mice inoculated with 10^5^ PFUs of JEV through the foot-pad route. Relative fold change in the (**A**) CD80, (B) CXCL11, (**C**) CLDN2, (**D**) IFNb, (**E**) IL1b, and (**F**) Il18 mRNA levels on 8^th^ dpi of JE virus-infected SIRT2^+/+^ or SIRT2^-/-^ mice brain (n=4 mice each group; The p-value was calculated using Student’s t-test).

**Figure S6**: **SIRT2 overexpression leads to a reduction in NF-**κ**B acetylation and lowers JEV-NS5 levels in cells infected with JEV and overexpressing SIRT2.** Immunoprecipitation of p65 at 30 hpi from JEV (1 MOI)- infected neuro2a cells treated with Ad-null or Ad-SIRT2, demonstrating p65 and SIRT2 interaction.

**Figure S7. The immune-blotting quantification of autophagy markers displaying the role of SIRT2 in the JEV induced autophagy.** (**A**) Immuno-blotting quantification displaying differences in autophagy markers at 30 hpi in the JE virus-infected neuro2a cells (n=3 independent experiments; duplicates were used in each experiment). The p-value was calculated using Student’s t-test. (**B**) Immune-blotting quantification displayed an increase in autophagy markers at 8 dpi in the JE virus-infected WT mice brain (n=6 mice in each group). The p-value was calculated using Student’s t-test. (**C**) Immuno-blotting quantification displaying differences in autophagy markers at 30 hpi in the JE virus-infected, control or SIRT2-KD neuro2a cells (n=3 independent experiments; duplicates were used in each experiment). The p-value was calculated using Student’s t-test. (**D**) Immuno-blotting quantification displaying change in autophagy markers at 30 hpi in the JE virus-infected, vehicle or AGK2-treated neuro2a cells (n=3 independent experiments; duplicates were used in each experiment). The p-value was calculated using Student’s t-test. (**E**) Immune-blotting quantification displaying differences in autophagy markers at 8 dpi in the SIRT2^+/+^ or SIRT2^-/-^JE virus-infected mice brain (n=6 mice in each group). The p-value was calculated using Student’s t-test. (**F**) Immune-blotting quantification displaying change in autophagy markers at 30 hpi in the JE virus-infected, Ad-null or Ad-SIRT2 treated neuro2a cells (n=3 independent experiments; duplicates were used in each experiment). Statistical analysis was performed using Student’s t-test.

**Figure S8. BAY-11, administered after the onset of JE clinical signs, significantly improved the survival of JE virus-infected mice.** (**A-C**) 10-day-old WT mice were infected through the foot-pad route with 10^5^ PFUs of JE virus, and vehicle or BAY-11 treatment started at the onset of JE clinical signs through the IP route every 24 hours. (**A**) The graph depicts the percentage of WT mice displaying paralysis in mock or JE virus-infected vehicles or BAY-11 treated mice (n=14 mice in each group). (**B**) The percentage of WT animals surviving in JE virus-infected vehicle or BAY-11 treated mice, in Kaplan-Meier survival percentage curve (n=14 mice each group). The p-value was calculated using the Log-rank (Mantel-Cox) test. (**C**) The graph represents JE virus PFUs in the JE virus-infected vehicle or BAY-11 treated WT mice brain (n=4 mice per dpi).

## Key resources used in the study

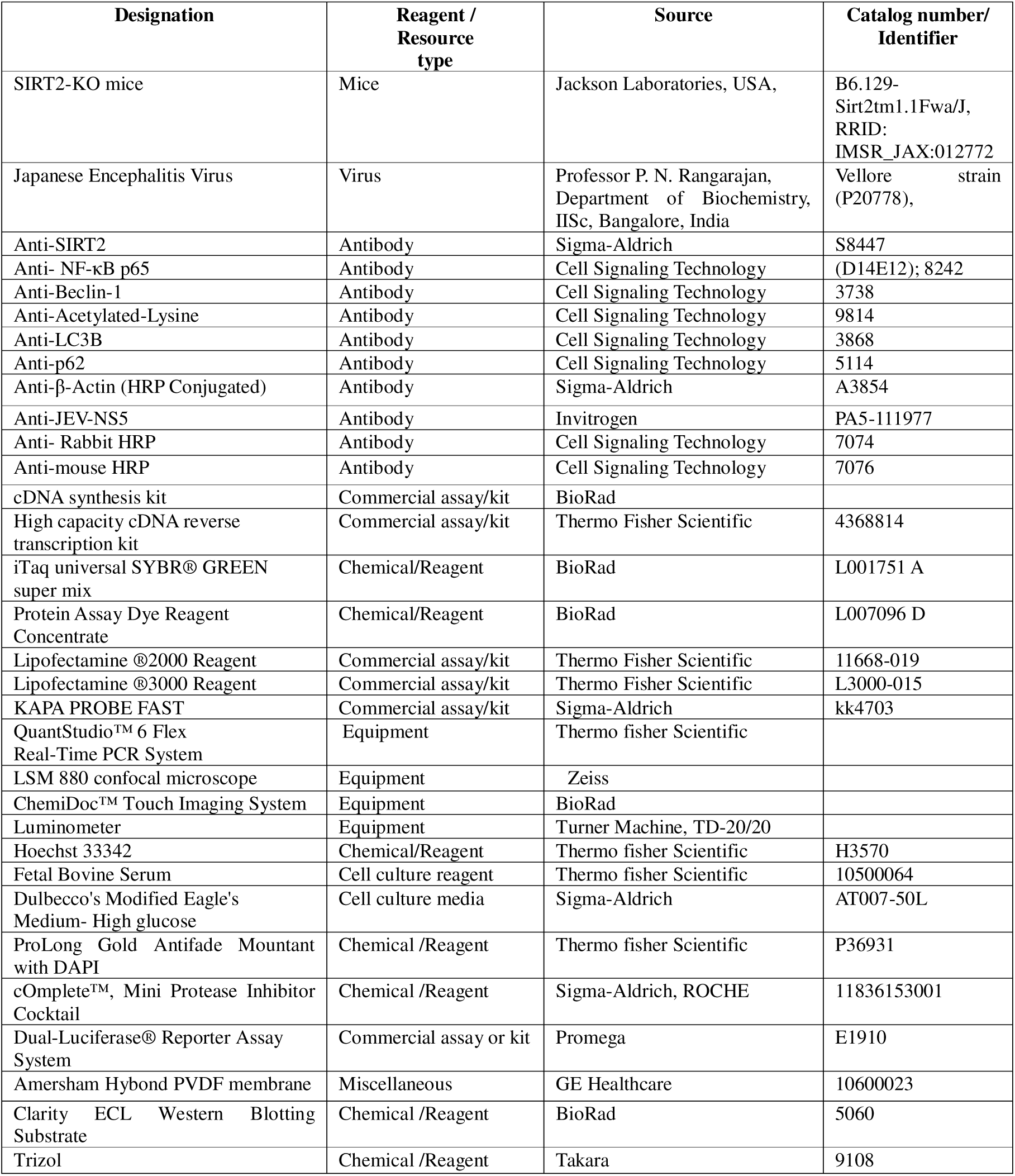
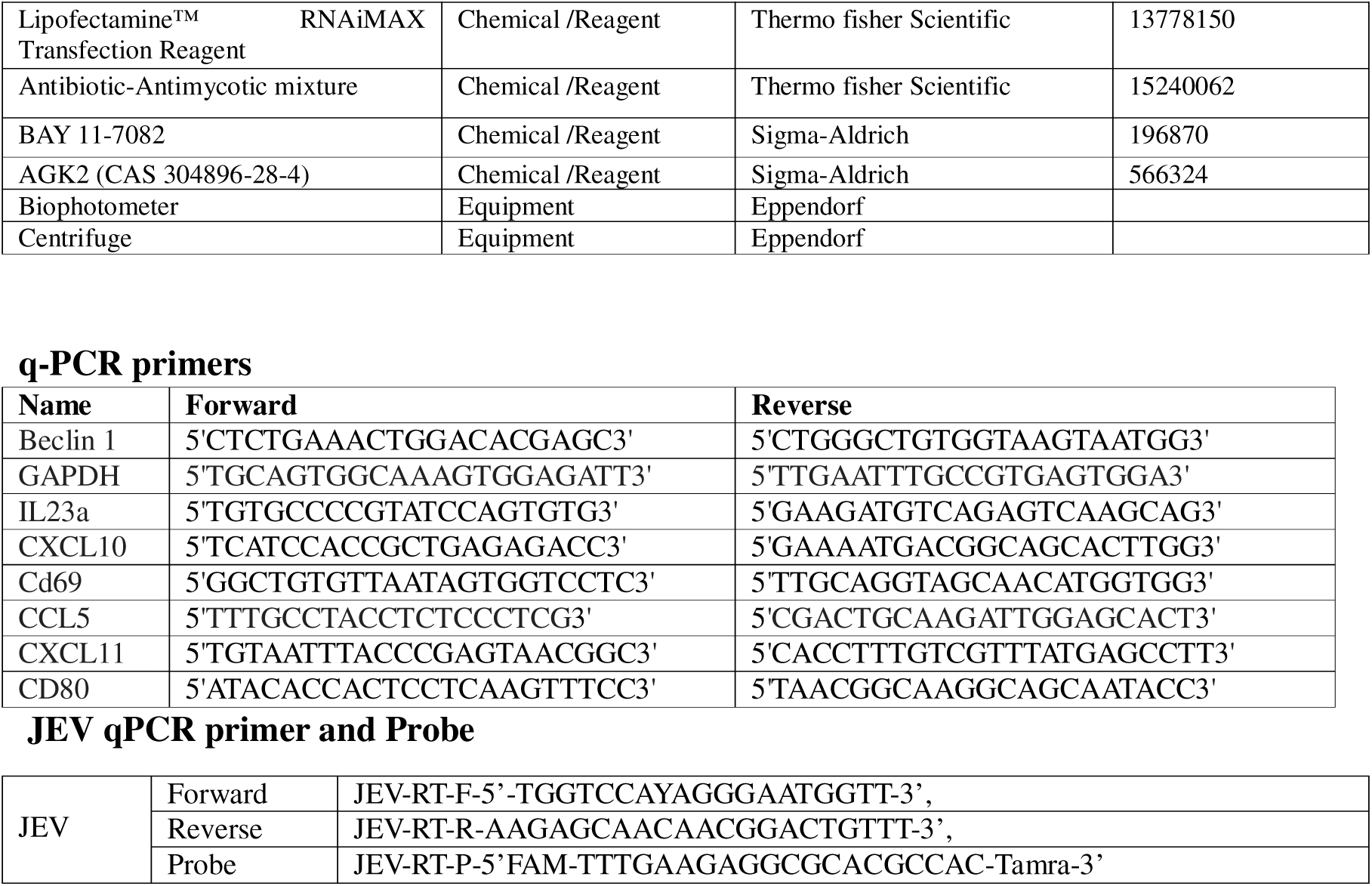

