## Supplementary figures and images for "SIRT2 protects against Japanese encephalitis virus infection in mice"

### Figure S1 New.tif

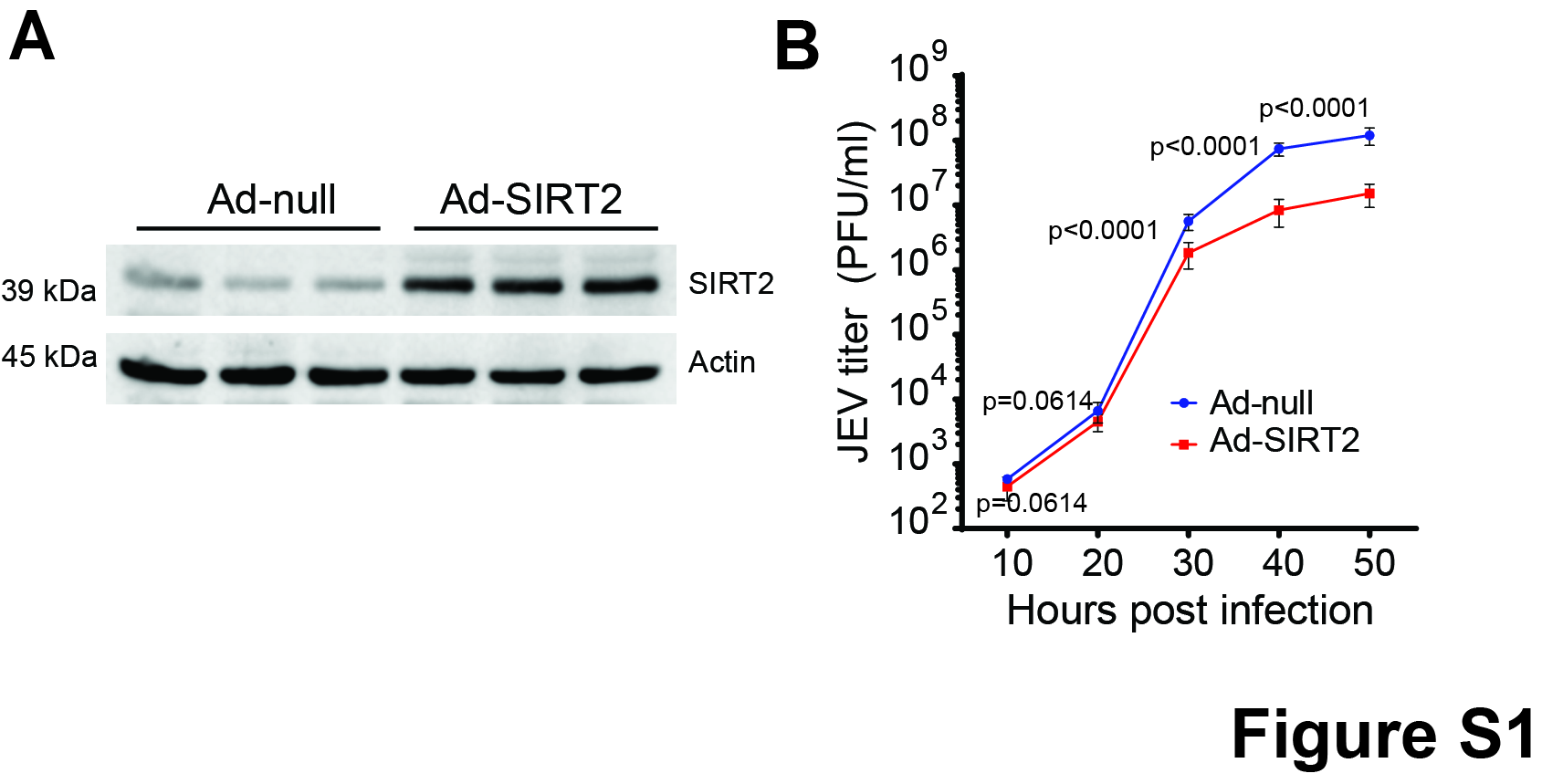

### Figure S2 new.tif

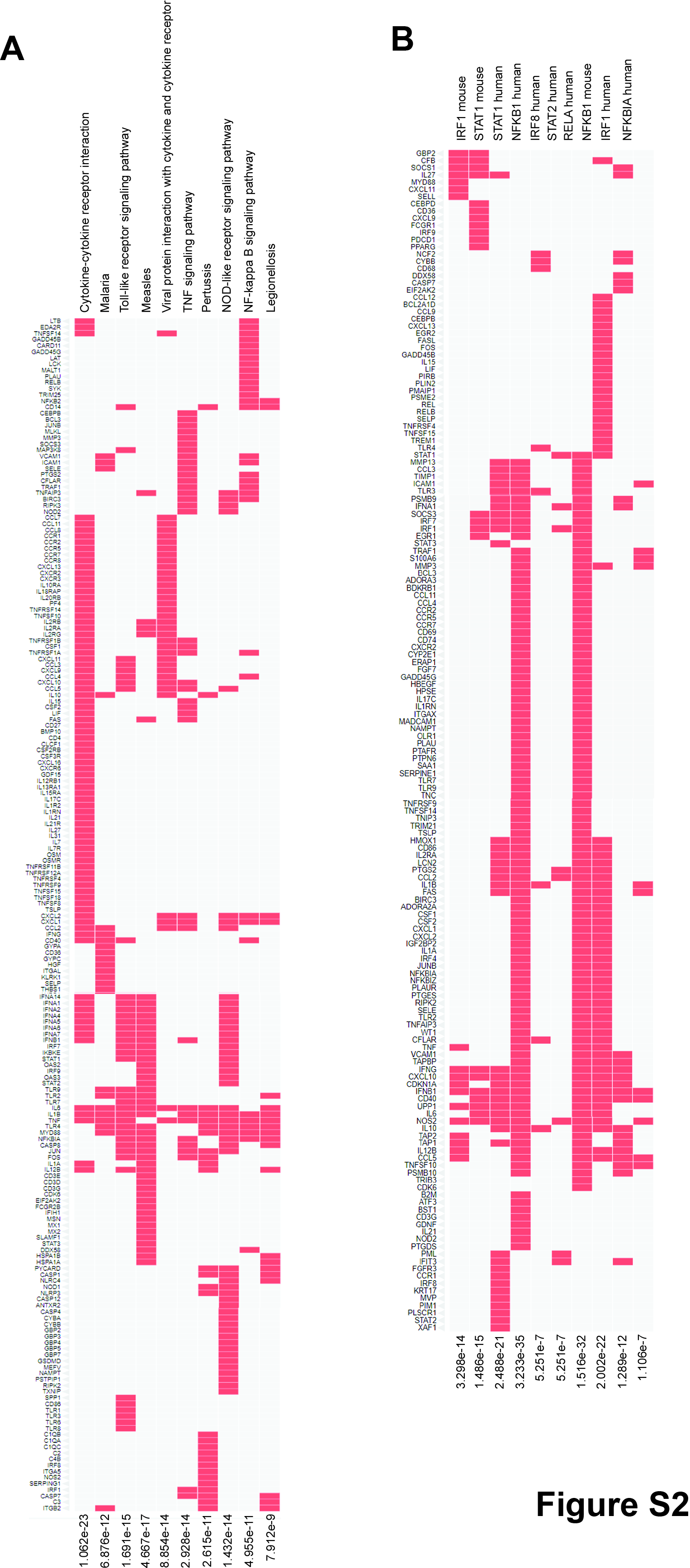

### Figure S3 new.tif

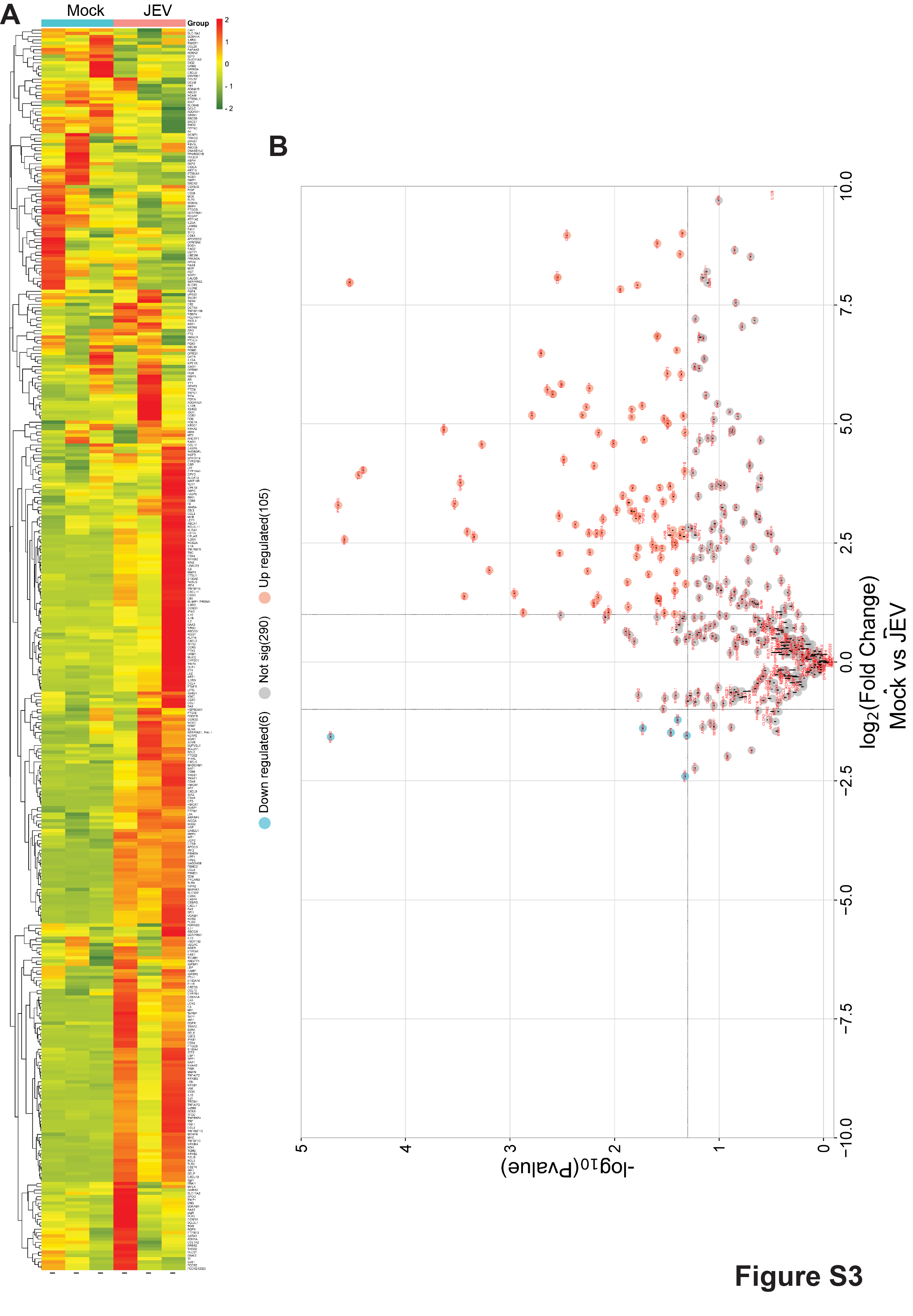

### Figure S4 new.tif

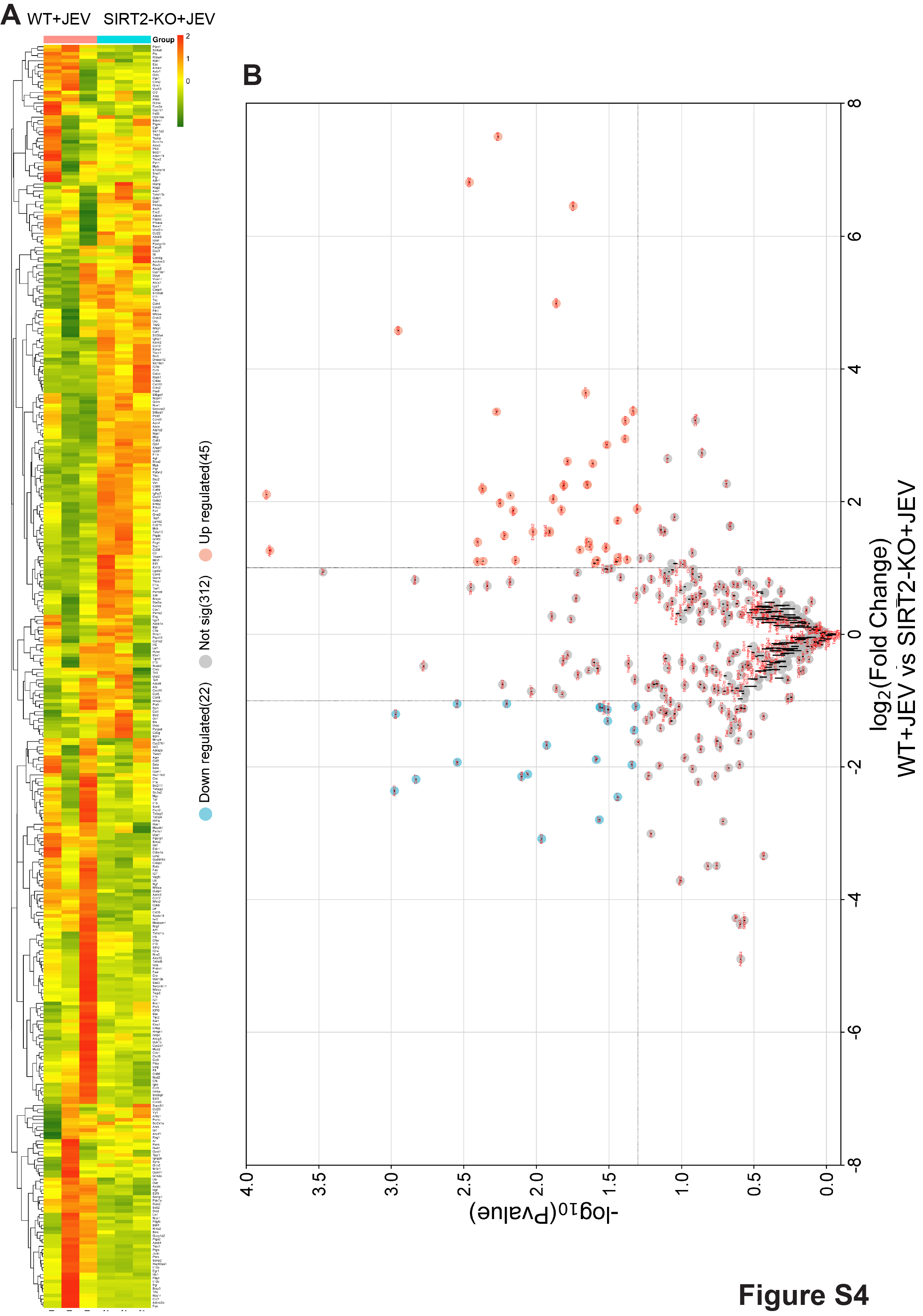

### Figure S5 Revision.tif

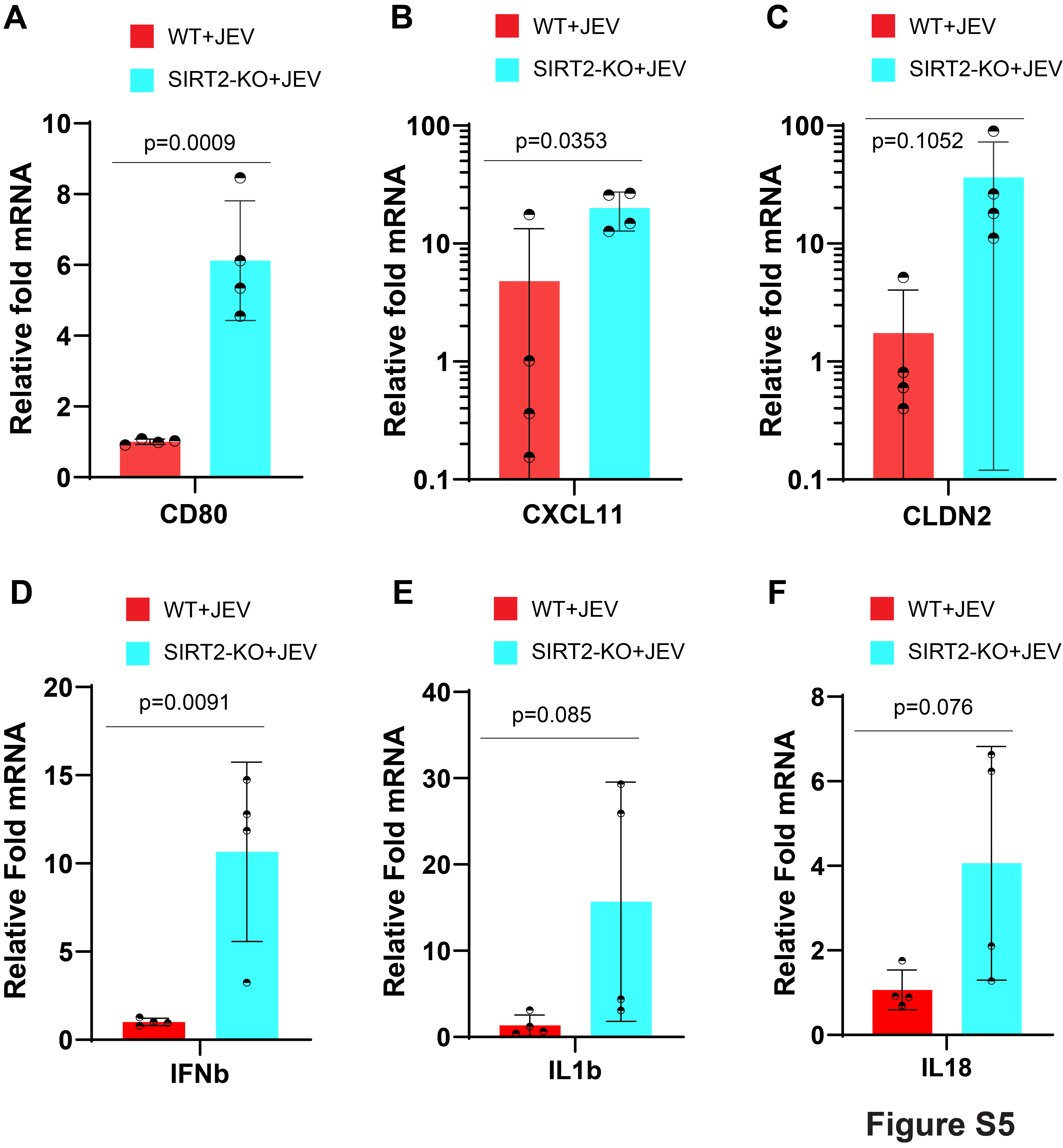

### Figure S6 Revision.tif

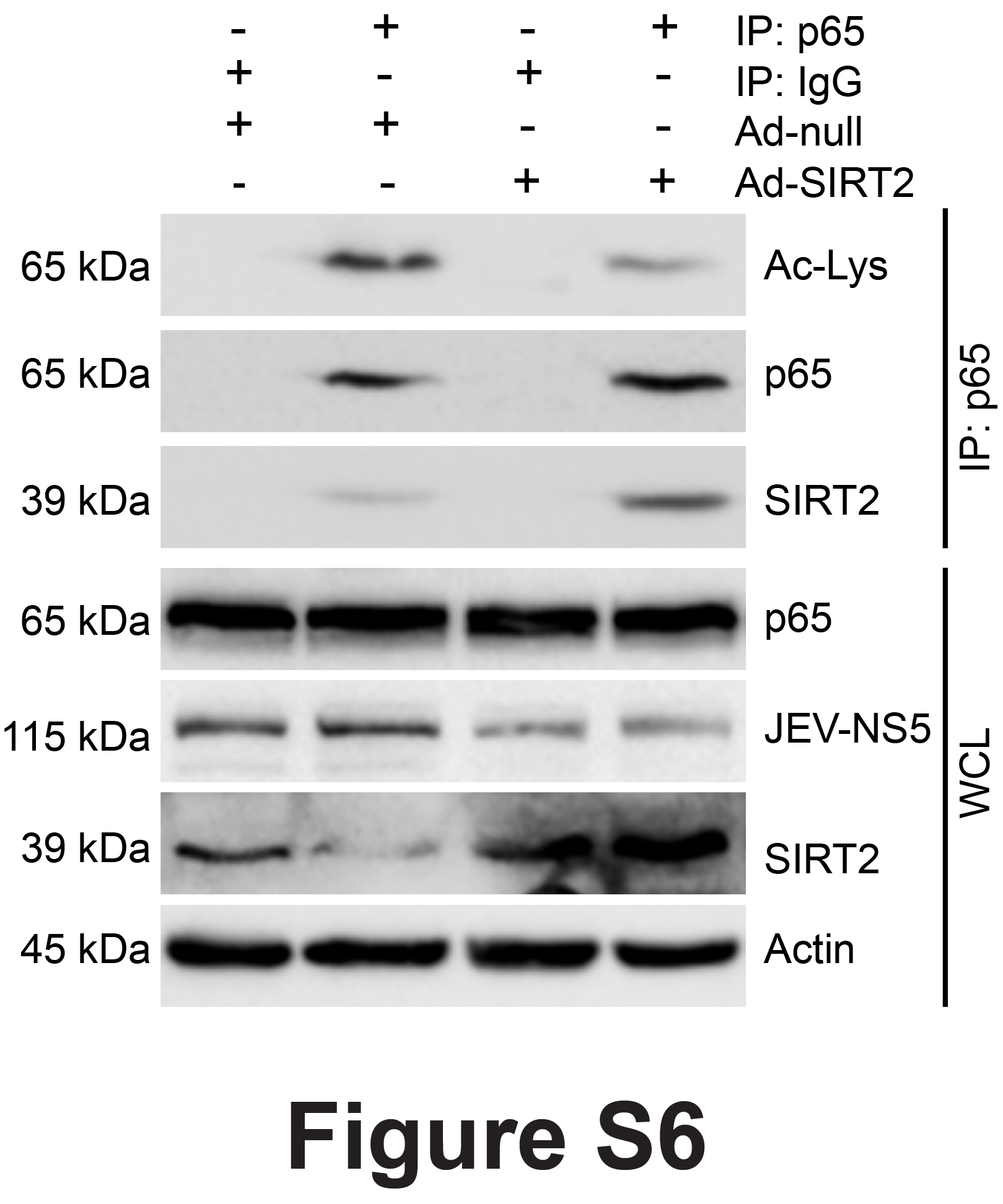

### Figure S7 revision.tif

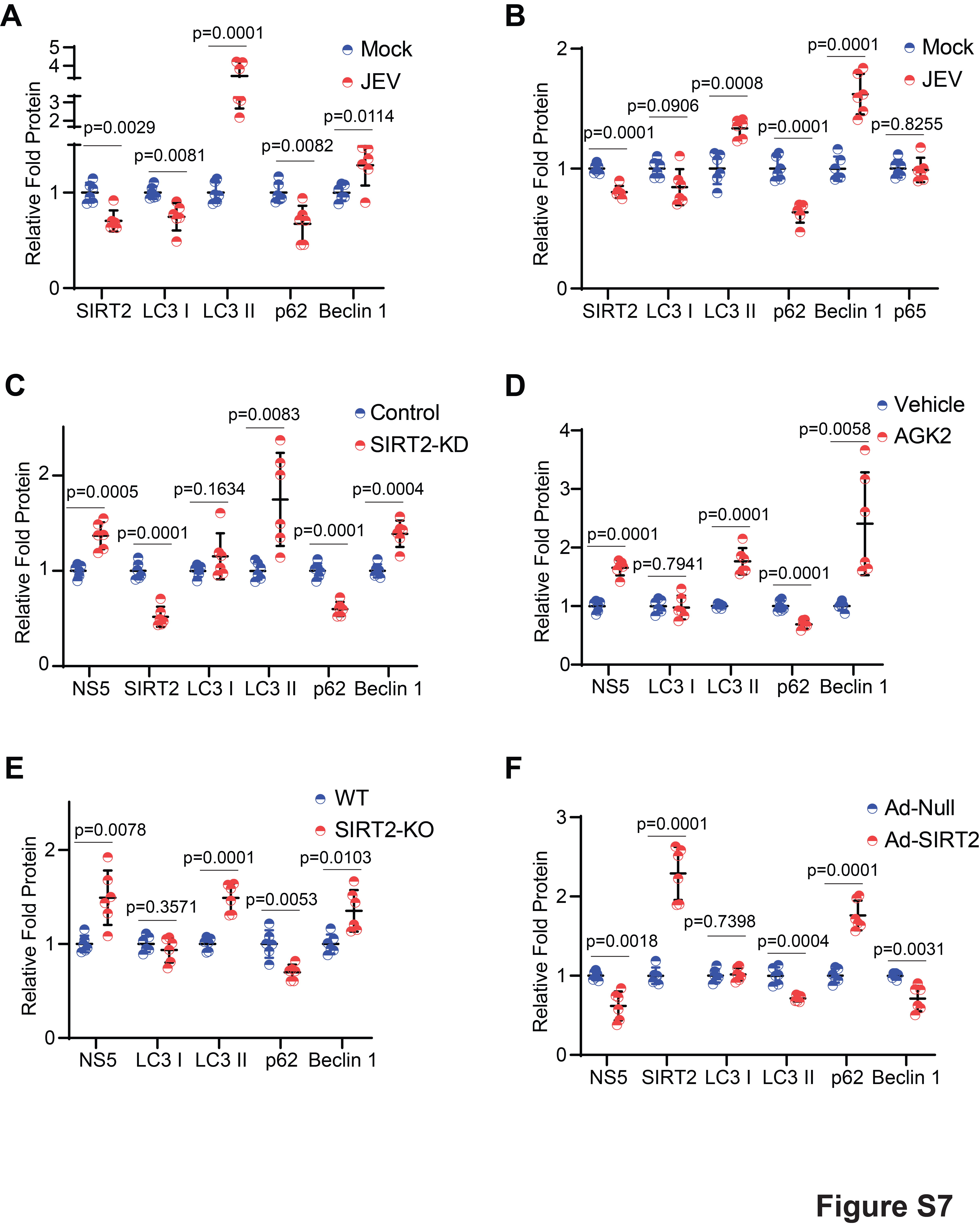

### Figure S8 revision.tif

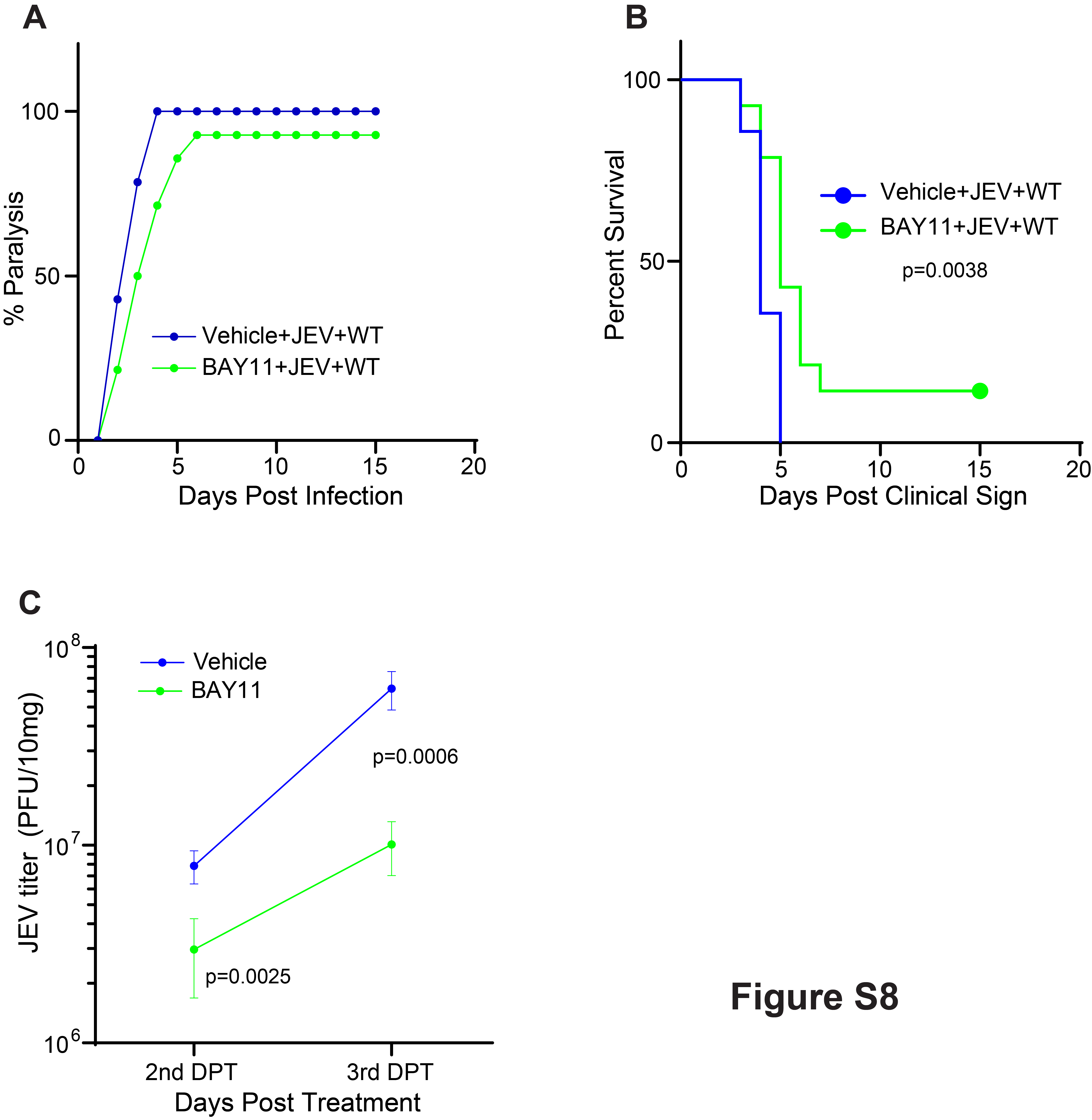
